# SARS-CoV-2 B.1.1.7 and B.1.351 variants of concern induce lethal disease in K18-hACE2 transgenic mice despite convalescent plasma therapy

**DOI:** 10.1101/2021.05.05.442784

**Authors:** Alexander M. Horspool, Chengjin Ye, Ting Y. Wong, Brynnan P. Russ, Katherine S. Lee, Michael T. Winters, Justin R. Bevere, Theodore Kieffer, Ivan Martinez, Julien Sourimant, Alexander Greninger, Richard K. Plemper, James Denvir, Holly A. Cyphert, Jordi Torrelles, Luis Martinez-Sobrido, F. Heath Damron

**Affiliations:** Department of Microbiology, Immunology, and Cell Biology, West Virginia University, Morgantown, WV, USA; Vaccine Development Center at West Virginia University Health Sciences Center, Morgantown, WV, USA; West Virginia University Cancer Institute, Morgantown, WV, USA; Department of Pathology, Anatomy and Laboratory Medicine, West Virginia University School of Medicine, Morgantown, WV, USA; Institute for Biomedical Sciences, Georgia State University, Atlanta, GA, USA; University of Washington, Department of Laboratory Medicine and Pathology, Seattle, Washington; Department of Biomedical Sciences, Marshall University, Huntington, WV. USA; Department of Biological Sciences, Marshall University, Huntington, WV, USA; Texas Biomedical Research Institute, San Antonio, TX, USA

**Keywords:** SARS-CoV-2, COVID-19, variants of concern, B.1.1.7, B.1.351, K18-hACE2 transgenic mouse, convalescent plasma

## Abstract

SARS-CoV-2 variants of concern (VoCs) are impacting responses to the COVID-19 pandemic. Here we present a comparison of the SARS-CoV-2 USA-WA1/2020 (WA-1) strain with B.1.1.7 and B.1.351 VoCs and identify significant differences in viral propagation *in vitro* and pathogenicity *in vivo* using K18-hACE2 transgenic mice. Passive immunization with plasma from an early pandemic SARS-CoV-2 patient resulted in significant differences in the outcome of VoC-infected mice. WA-1-infected mice were protected by plasma, B.1.1.7-infected mice were partially protected, and B.1.351-infected mice were not protected. Serological correlates of disease were different between VoC-infected mice, with B.1.351 triggering significantly altered cytokine profiles than other strains. In this study, we defined infectivity and immune responses triggered by VoCs and observed that early 2020 SARS-CoV-2 human immune plasma was insufficient to protect against challenge with B.1.1.7 and B.1.351 in the mouse model.

## INTRODUCTION

The evolution of Severe Acute Respiratory Syndrome CoV-2 (SARS-CoV-2) VoCs has been a source of escalating epidemiological alarm in the currently ongoing coronavirus disease 2019 (COVID-19) pandemic. Mutants of SARS-CoV-2 have emerged and are thought to be more infectious and more lethal than the early 2020 original Wuhan-Hu-1 or USA-WA1/2020 (WA-1) strains (Challen et al., 2021; Korber et al., 2020; Toyoshima et al., 2020). The VoC B.1.1.7, first identified in the United Kingdom (Rambaut et al., 2020), and B.1.351, first identified in South Africa (Tegally et al., 2020), are two emerging SARS-CoV-2 VoCs that are rapidly spreading around the world and exhibit high levels of infectivity and therapeutic resistance (Challen et al., 2021; Chen et al., 2021; Davies et al., 2021; Galloway et al., 2021; 2021b, 2021a; Wang et al., 2021). Both VoCs harbor significant evolution in the receptor binding domain (RBD) of the spike (S) viral glycoprotein (Rambaut et al., 2020; Tegally et al., 2020) that are predicted to impact binding to the human angiotensin converting enzyme 2 (hACE2) viral receptor and enhance viral entry to host cells (Bozdaganyan et al., 2021; Laffeber et al., 2021; Ozono et al., 2021; Shah et al., 2020; Tian et al., 2021). In particular, B.1.1.7 contains the D614G, and N501Y, mutations in the SARS-CoV-2 S RBD which are theorized to increase the ability of the virus to bind to hACE2 (Ozono et al., 2021; Tian et al., 2021). B.1.351 possesses these key mutations in the S RBD, in addition to the K417N mutation E484K mutation which are not directly implicated in altered viral transmission and hACE2 binding (Laffeber et al., 2021; Zhou et al., 2021). The culmination of high infectivity, therapeutic resistance, and key changes in the viral genome suggests that these VoCs may have an impact on pathogenicity in animal models of SARS-CoV-2. This could have an impact on evaluating SARS-CoV-2 pathogenesis as well as prophylactic (vaccines) and therapeutics (antivirals).

The K18-hACE2 transgenic mouse model (McCray et al., 2007) of SARS-CoV-2 infection was established by Perlman and McCray among others in 2020 (Moreau et al., 2020; Oladunni et al., 2020; Winkler et al., 2020). K18-hACE2 transgenic mice infected with SARS-CoV-2 exhibit significant morbidity and mortality, viral tropism of the respiratory and central nervous systems, elevated systemic chemokine and cytokine levels, significant tissue pathologies, and altered gross clinical measures (Oladunni et al., 2020; Winkler et al., 2020; Yinda et al., 2021; Zheng et al., 2021). The generation of this mouse model has led to numerous studies of SARS-CoV-2 infection for a variety of purposes including understanding SARS-CoV-2 related immunity, and therapeutic / vaccine testing (Hassan et al., 2020; Kumari et al., 2020; Liu et al., 2021; Moreau et al., 2020; Pandey et al., 2020; Sarkar and Guha, 2020; Silvas et al., 2021). As the world experiences an increase in the number of SARS-CoV-2 VoCs, it is imperative to adapt existing preclinical animal infection models to these newly emerging VoC. Specifically, it is critical to understand if the K18-hACE2 transgenic mouse model first, is useful for studying SARS-CoV-2 VoC infection dynamics and second, if it exhibits any differences after challenge with newly emerged SARS-CoV-2 VoCs. An investigation of these key points will provide context for studies important for developing new therapeutics and prophylactics as the COVID-19 pandemic continues and as new VoCs emerge.

Clinical studies of therapeutics and vaccines for COVID-19 have been complicated by the rise of SARS-CoV-2 VoCs. Therapeutic escape by these mutants is documented (Wang et al., 2021) and requires the development of novel treatment options as well as re-evaluation of existing ones. One of the first treatment options for COVID-19 was infusion of convalescent plasma (CP). CP exhibited some beneficial effects early in the pandemic for critically ill patients (Bloch, 2020; Bloch et al., 2020; Chen et al., 2020), but its utility has recently been called into question (Casadevall et al., 2021; Cele et al., 2021; NIH, 2021; Zhao and He, 2020). There are speculations as to the reasons behind this, including that neutralizing antibodies (nAbs) generated against the original Wuhan-Hu-1 or WA-1 SARS-CoV-2 S RBD may have different affinity to the new VoCs. As many therapeutics and vaccines have focused on the RBD or S protein of SARS-CoV-2 Wuhan or WA-1, it was of interest to determine whether early pandemic convalescent plasma containing neutralizing antibodies against WA-1 SARS-CoV-2 S RBD protects against these VoCs with mutations in their RBD. Thus, the overall focus of this study was to observe the effects of SARS-CoV-2 VoCs on the K18-hACE2 transgenic mouse model of COVID-19, and determine whether theses VoCs can evade an early COVID-19 pandemic therapeutic: CP treatment.

## METHODS

### Ethics and biosafety

Human plasma used in this study was obtained under WVU IRB no. 2004976401. Experiments with live SARS-CoV-2 virus were conducted in Biosafety Level 3 (BSL-3) Texas Biomedical Research Institute (TBRI IBC BSC20-004) or West Virginia University (WVU IBC 20-09-03). All ABSL-3 animal experiments were conducted under West Virginia University (WVU) ACUC protocol no. 2009036460.

### Viral growth and in vitro analysis of SARS-CoV-2 replication

SARS-CoV-2 USA-WA-1/2020 (NR-52281) (WA-1), B.1.1.7 (NR-54000), and B.1.351 (NR-54008) strains were obtained from BEI Resources and propagated in Vero E6 cells (ATCC-CRL-1586) as previously described (Case et al., 2020; Oladunni et al., 2020). Vero E6 cells for viral titrations (6-well plate, 10^6^ cells/well) were infected with serial dilutions of SARS-CoV-2 WA-1, B.1.1.7 or B.1.351 VoCs. At 72 hours post-infection, cells were fixed overnight with 10% formalin (Sigma HT501128-4L), permeabilized and immunostained with 1µg/mL of a SARS-CoV cross-reactive nucleocapsid (N) protein antibody 1C7C7, kindly provided by Dr. Thomas Moran at the Icahn School of Medicine at Mount Sinai. For viral growth kinetics, Vero E6 cells (6-well plate, 10^6^ cells/well, triplicates) were infected (MOI 0.01) with SARS-CoV-2 WA-1, B.1.1.7 or B.1.351. At the indicated times after infection (12, 24, 48 and 72 hours), tissue culture samples were collected and titrated by plaque assay as described previously (Oladunni et al., 2020).

### Sequencing of SARS-CoV-2 VoCs

SARS-CoV-2 viral RNA from all stocks used for *in vitro* analyses was deep sequenced according to the method described previously (Ye et al., 2020a). Briefly, we generated libraries using KAPA RNA HyperPrep Kit (Roche KK8541) with a 45 min adapter ligation incubation including 6-cycle of PCR with 100 ng RNA and 7 mM adapter concentration. Samples were sequenced on an Illumina Hiseq X machine. Raw reads were quality filtered using Trimmomatic v0.39 (Bolger et al., 2014) and mapped to a SARS-CoV-2 reference genome (Genbank Accession No. MN985325) with Bowtie2 v2.4.1 (Langmead and Salzberg, 2012). Genome coverage was quantified with MosDepth. version 0.2.6 (Pedersen and Quinlan, 2018). We genotyped each sample for low frequency VoCs with LoFreq* v2.1.3.1 (Wilm et al., 2012) and filtered sites with allele frequencies less than 20%. SARS-CoV-2 viral RNA from stocks used for K18-hACE2 transgenic mice infection was deep sequenced and reads were aligned to the MN908947.3 reference genome using BWA version 0.7.17 (Li and Durbin, 2009) and trimmed for base-calling quality using iVar version 1.3.1 (Grubaugh et al., 2019) with default parameters. Consensus sequence and individual mutations relative to the reference genome were determined using iVar, with a minimum allele frequency of 30% used as a threshold for calling a mutation. Coverage was computed using samtools mpileup version 1.11 (Li et al., 2009). Lineage was confirmed using pangolin version 2.3.5 and pangoLEARN version 2021-03-16 (O’Toole et al.). Authentication of the B.1.351 stock was performed using metagenomic sequencing as described previously (Addetia et al., 2020; Greninger et al., 2017). Viral RNA was treated with Turbo DNase I (Thermo Fisher). cDNA was generated from random hexamers using SuperScript III reverse transcriptase, second strand was generated using Sequenase 2.0, and cleaned using 0.8× Ampure XP beads purification on a SciClone IQ (Perkin Elmer). Sequencing libraries were generated using two-fifths volumes of Nextera XT on ds-cDNA with 18 cycles of PCR amplification. Libraries were cleaned using 0.8×Ampure XP beads and pooled equimolarly before sequencing on an Illumina NovaSeq (1×100bp run). Raw fastq reads were trimmed using cutadapt (-q 20) (Martin). To interrogate potential resistance alleles, reference-based mapping to NC_045512.2 was carried out using our modified Longitudinal Analysis of Viral Alleles (LAVA -https://github.com/michellejlin/lava) (Jin et al., 2019) pipeline. LAVA constructs a candidate reference genome from early passage virus using bwa (Li and Durbin, 2009), removes PCR duplicates with Picard, calls variants with VarScan (Koboldt et al., 2009, 2012), and converts these changes into amino acid changes with Annovar (Wang et al., 2010). The genome sequence for strain B.1.351 is accession number MZ065365 and SRA BioProject PRJNA726258.

### Infection of K18-hACE2 transgenic mice with SARS-CoV-2 VoCs and treatment with human Plasma

SARS-CoV-2 VoCs were thawed from −80°C and diluted in infection medium (Dulbecco’s Modified Eagle Medium 4/.5g/L glucose + 2% fetal bovine serum + 1% HEPES + 1% penicillin/streptomycin at 10,000 units/µg/mL) to a concentration of 10^6^ plaque forming units (pfu) /mL in the WVU BSL-3 high-containment facility. Male and female (Figures 2-3) or male (Figures 4-7) eight-week-old B6.Cg-Tg(K18-hACE2)2Prlmn/J mice (Jackson Laboratory 034860) were anesthetized with a single intraperitoneal dose of ketamine (Patterson Veterinary 07-803-6637, 80 mg/kg) + xylazine (Patterson Veterinary 07-808-1947, 8.3 mg/kg) and the 50µL infectious dose was administered with a pipette intranasally, 25µL per nare. 500µL of convalescent plasma (CP) or healthy human sera (HHS) with known anti-SARS-CoV-2 IgGs and nAbs (Supplementary Figure 1) were administered intraperitoneally at this time. Convalescent human plasma was obtained from a single individual with PCR-confirmed SARS-CoV-2 infection in March 2020 via WVU IRB no. 2004976401. Mice were monitored until awake and alert.

**Figure 1.**
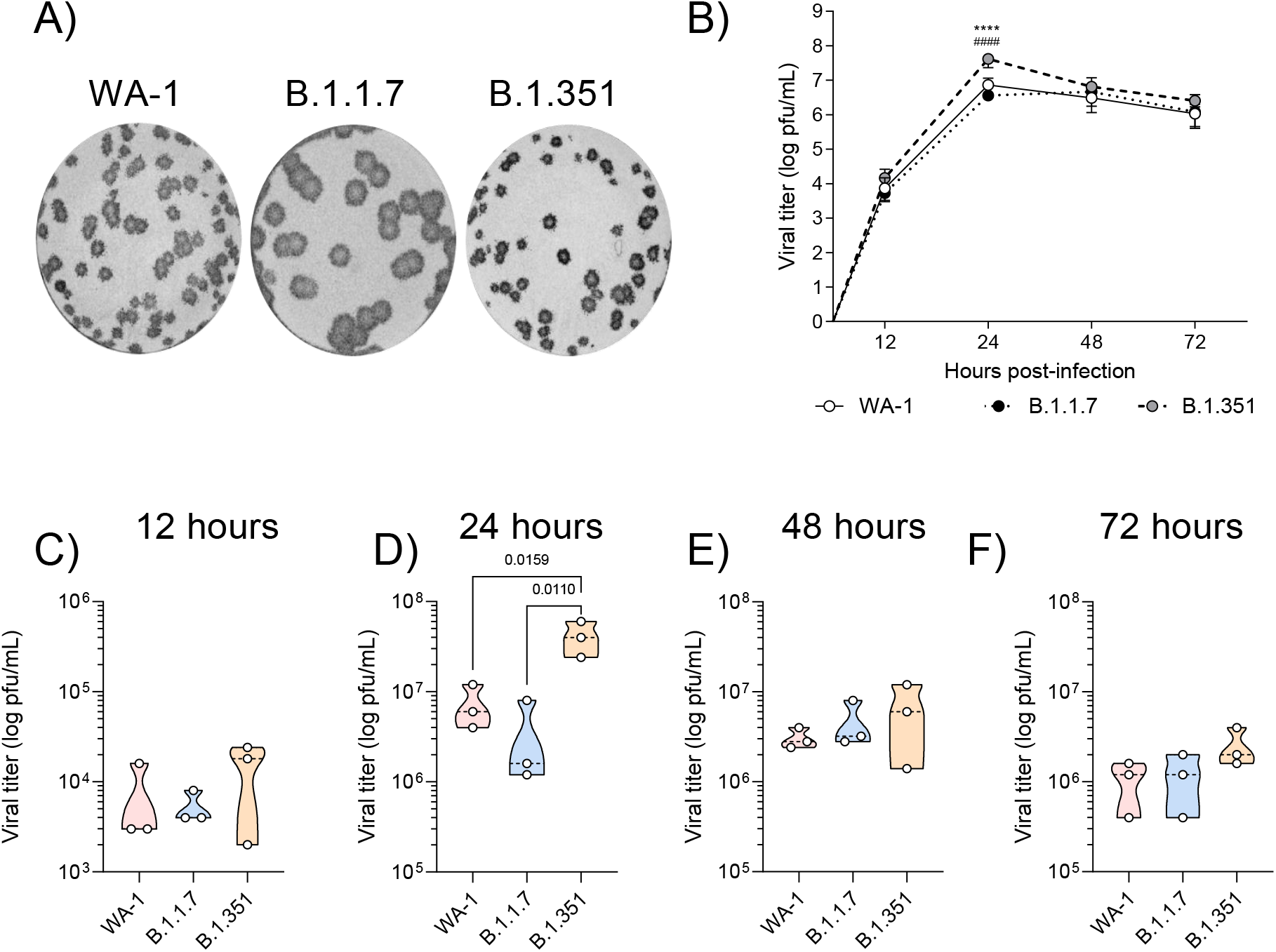
Characterization of SARS-CoV-2 variants *in vitro*: (A) Plaque morphology of SARS-CoV-2 WA-1, B.1.1.7 or B.1.351 infected VeroE6 cells. (B) Quantification of viral replication of SARS-CoV-2 variants in VeroE6 cells over time was quantified. Comparison of viral titers in cell culture at 12 (C), 24 (D), 48 (E), and 72 (F) hours post-infection. Statistical analysis of viral replication was completed by two-way ANOVA followed by Tukey’s multiple comparison test, or RM ANOVA followed by Tukey’s multiple comparison test. **** = *P* < 0.0001 relative to WA-1, ^####^= *P* < 0.0001 relative to B.1.1.7.

**Figure 2.**
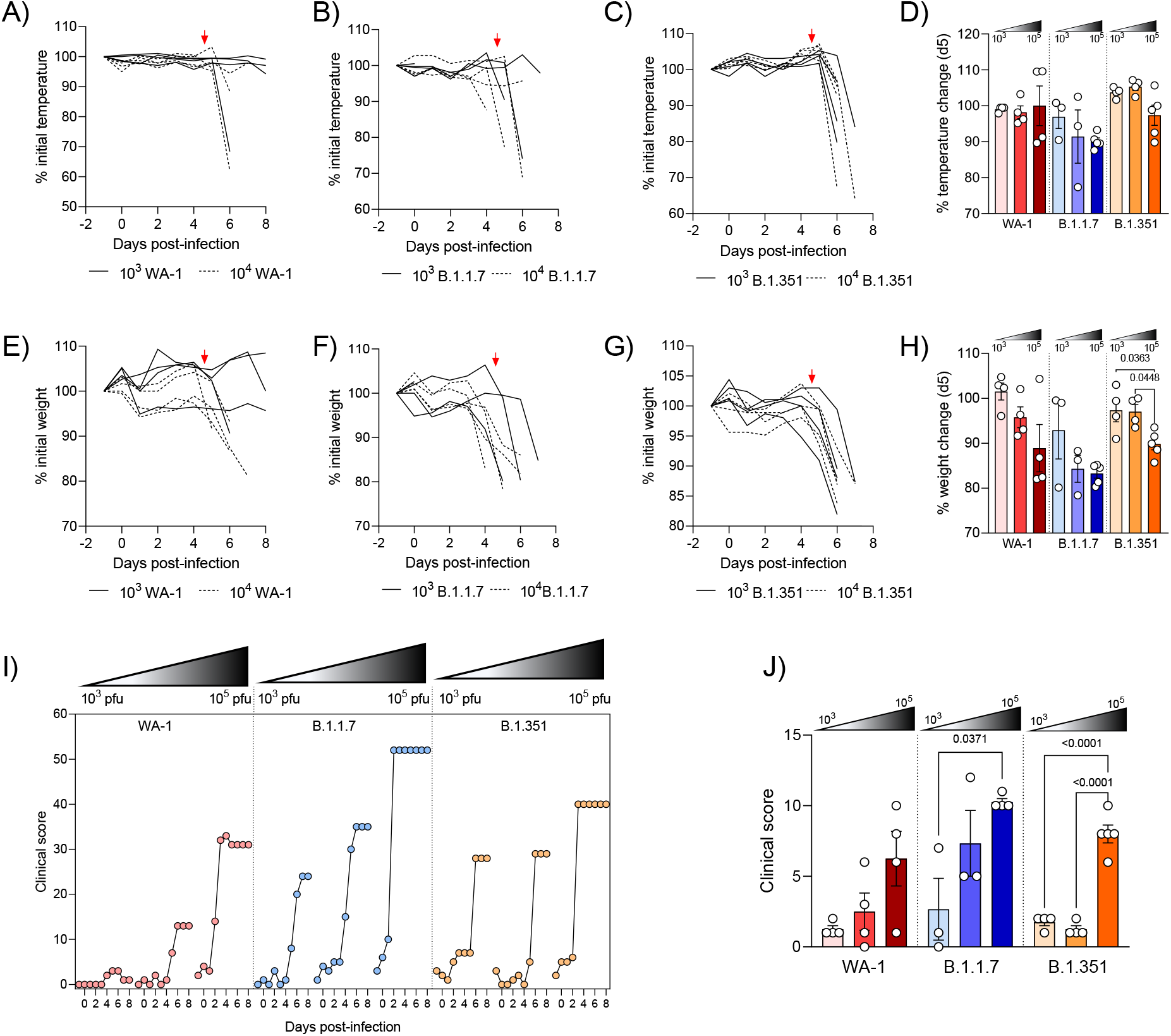
Establishing clinical endpoints of SARS-CoV-2 VoC infection in K18-hACE2 transgenic mice: Mice were infected with 10^3^, 10^4^ or 10^5^ pfu of SARS-CoV-2 VoCs and were monitored for temperature (A-C), body weight (E-G), and clinical score (I-J) changes over infection. Temperature (D), body weight (H), and clinical scores (J) on day 5 post-infection were assessed. If mice were deceased at five days post-infection, their clinical data at time of euthanasia is presented. Arrows indicate day 5 (A-H). Statistical significance was assessed by one-way ANOVA followed by Tukey’s multiple comparison test. n > 3 subjects per group.

**Figure 3.**
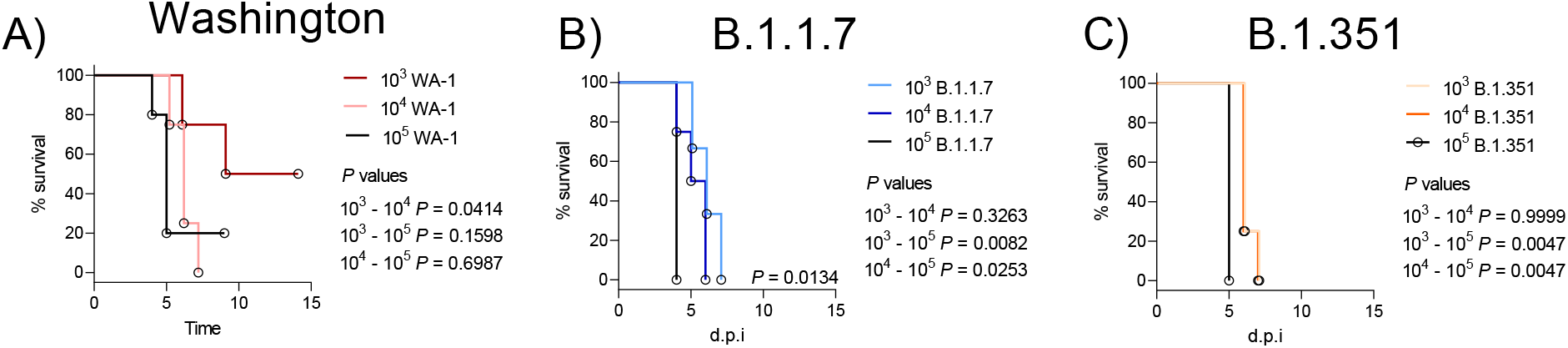
Impact of dose on survival of K18-hACE2 transgenic mice infected with SARS-CoV-2 VoCs: Kaplan-Meyer survival curves of mice infected with WA-1 (A), B.1.1.7 (B), or B.1.351 (C) VoCs at 10^3^, 10^4^, and 10^5^ pfu doses. Statistical significance was assessed by Mantel-Cox tests. n = 3-5 subjects per group. *P* values for significant differences are reported.

**Figure 4.**
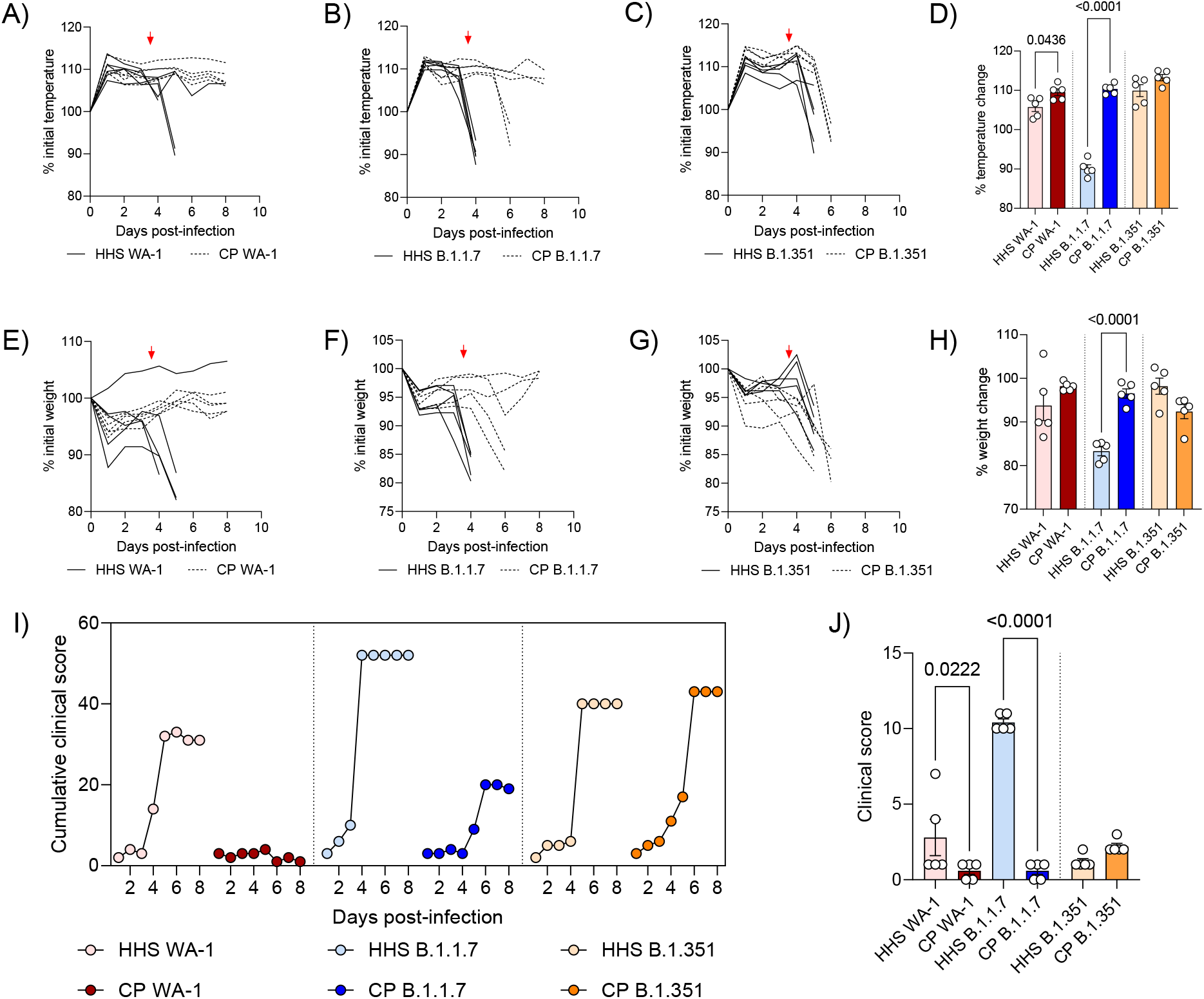
Effect of convalescent plasma treatment on SARS-CoV-2 VoC infection in K18-hACE2 transgenic mice: Mice infected with 10^5^ pfu SARS-CoV-2 VoCs were treated intraperitoneally with 500µL HHS or CP and monitored for temperature (A-C), body weight (E-G) and clinical score (I) over the course of infection. Temperatures (D), body weights (H), and clinical scores (J) on day 4 post-infection were assessed. Arrows indicate day 4 (A-H). Statistical significance was assessed by one-way ANOVA followed by Tukey’s multiple comparison test. N > 3 subjects per group. *P* values for significant differences are reported.

### Clinical scoring of SARS-CoV-2 infected mice

Mice were scored daily on a scale encompassing appearance (score of 0-2), eye health (score of 0-2), respiration (score of 0-2), activity (score of 0-3), weight loss (score of 0-5), and hypothermia (0-2) (Supplementary Figure 1). Appearance included visual identification of a combination of mild to severe piloerection (0-2) or lack of grooming (0-2). Eye health scores were defined by observation of squinting (1), prolonged eye closure not related to sleep (2), or eye discharge (0-2) depending on severity. The maximal combined score for eye health was 2. Respiration (assessed visually) outside the range of 80-240 breaths per minute required mandatory euthanasia and scored as 2. Respiration that was abnormal in regularity was scored as 1. Activity was scored as slow (1), immobile (2), or collapsed and immobile (3). Weight loss was scored as 0-5% (0), 5-10% (1), 10-15% (2) 15-20% (3), >20% (4-5). All mice with weight loss greater than 20% were humanely euthanized. Hypothermia was assessed and scored as not-present (>36.4°C, 0), developing (36.4°C – 35.0°C, 1) or present (<35.0°C, 2) (Supplementary Figure 1).

### Euthanasia and necropsy of SARS-CoV-2 infected mice

Euthanasia was conducted by administering 200 µL of pentobarbital (Patterson Veterinary 07-805-9296, 390 mg/kg diluted in 0.9% sterile NaCl) and cardiac puncture. Blood was aliquoted into gold serum separator tubes (BD 365967) and centrifugated at 15,000 x *g* for 5 min. Serum was removed and stored in 1.5 mL tubes at −80°C until needed. Lungs were removed from animals and the right lobes of the lung were homogenized in 1mL of PBS in Miltentyi C tubes (Miltenyi Biotec 130-096-334) using the m_lung_02 program on a Miltenyi gentleMACS tissue dissociator. An aliquot of each lung homogenate (300µL) was added to 100µL of TRIReagent (Zymo Research R2050-1-200) and stored at −80°C. Remaining homogenates (300µL) were spun down at 15,000 x *g* and the supernatants collected. Pellets were frozen at −80°C until use. Brain tissue was removed from animals and split down the mid-line. The right brain was added to 1mL of PBS in Miltenyi C tubes and homogenized using the m_lung_02 program. An aliquot of each homogenate (500µL) was added 167µL aliquots of TRIReagent and stored at −80°C until use. Remaining homogenates were frozen at −80°C until use. To inactivate virus from tissue samples, 1% v/v Triton X-100 (Sigma-Aldrich T8787) (Winkler et al., 2020) was added to each sample and incubated for 1 hour at room temperature. Inactivated samples were then removed from the ABSL-3 facility.

### Evaluating viral copy number in SARS-CoV-2 infected tissues

RNA from homogenized virus-inactivated lung and brain tissues of SARS-CoV-2 infected animals was extracted using the Direct-zol RNA MiniPrep Kit (Zymo Research R2051) following the manufacturer’s instructions. RT-PCR and qPCR were performed by generating a master mix of: 10µL of TaqMan RT-PCR Mix from the Applied Biosystems TaqMan RNA to CT One Step Kit (Thermo-Fisher Scientific 4392938), 900nM (1.8µL) of (ATGCTGCAATCGTGCTACAA) forward nucleocapsid primer (Winkler et al., 2020), 900nM (1.8µL) of (GACTGCCGCCTCTGCTC) reverse nucleocapsid primer (Winkler et al., 2020), 250nM (0.5µL) of TaqMan probe (56-FAM/TCAAGGAAC/ZEN/AACATTGCCAA/3IABkFQ), 0.5µL of TaqMan RT enzyme from the Applied Biosystems TaqMan RNA to CT One Step Kit (Thermo-Fisher Scientific 4392938), 100ng of RNA, and RNAse/DNAse free water to make a 20µL total reaction volume. Samples were run in triplicate in Microamp Optical 96-well Fast Reaction Plates (Thermo-Fisher Scientific 4306737) through the following protocol: reverse transcription at 48°C for 15 minutes, activation of AmpliTaq Gold DNA polymerase at 95°C for 10 minutes, and 50 cycles of 95°C denaturing for 10 seconds followed by 60°C annealing for 60 seconds. Samples were run on an Applied Biosystems StepOnePlus Real-Time PCR System. Samples with undetectable virus were assigned a value of 1. C_T_ values and copy numbers were calculated and analyzed in Microsoft Excel and GraphPad Prism v9.0.0.

### Assessment of human IgGs against WA-1 SARS-CoV-2 S RBD and N

Human IgGs against WA-1 SARS-CoV-2 S RBD and N were quantified using ELISA as described previously (Horspool et al., 2021). Briefly, WA-1 S RBD (2µg/mL) or N (1µg/mL) proteins were coated on plates and blocked with 3% milk in 0.1% Tween 20 +PBS (PBS-T). Plates were washed three times with PBS-T (200µL) and virus inactivated samples (25µL) from infected mice were added to 100µL of sample buffer (1% milk + 0.1% Tween 20 diluted in PBS) and serially diluted (5-fold) down the plates. The final row was left with 100µL of sample buffer as a negative control. Plates were incubated for 10 minutes at room temperature shaking at 60rpm and subsequently washed four times with PBS-T (200µL). Secondary antibody (100µL 1:500 anti-human IgG HRP, Invitrogen 31410) was added and plates were incubated for 10 minutes at room temperature shaking at 60rpm. After incubation, plates were washed five times with PBS-T (200µL) and SigmaFAST OPD (Sigma-Aldrich P9187, 100µL) was added to each well of the plate. OPD development was stopped with 25µL of 3M hydrochloric acid and plates were read at an absorbance of 492nm on a Synergy H1 plate-reader. Area under the curve analysis was completed in GraphPad Prism. Human samples used as a comparison in Supplementary Figure 4 were obtained via WVU IRB no. 2004976401 as described previously (Horspool et al., 2021).

### Quantification of nAbs against WA-1 SARS-CoV-2 S RBD

An assay to assess nAb levels was developed using Luminex bead and Magpix technologies. SARS-CoV-2 S RBD (1µg) produced at WVU as described previously(Horspool et al., 2021) was conjugated to Luminex MagPlex® Microspheres (MC10012-YY) using the Luminex xMAP antibody coupling kit (Luminex 40-50016) per the manufacturer’s instructions. Conjugated beads (50µL containing 2000 beads suspended in 1x PBS-TBN (Phosphate buffered saline + 0.1% bovine serum albumin + 0.02% Tween 20 +0.05% sodium azide) diluted in de-ionized water from 5x PBS-TBN (Teknova P0211) were loaded into black non-binding Greiner 96-well plates (Greiner 655900). Human plasma/serum samples (25µL) were added into 100µL of PBS in the first row of a second black non-binding plate. Samples were serially diluted (5-fold dilution in PBS) down the plate. The final row contained PBS as a negative control. Diluted serum samples (50µL) were added to the 96-well plate containing the beads, creating a total reaction volume of 50µL beads (2000 beads), and 50µL diluted serum. The plates were covered with foil and shaken at 700rpm for 1 hour at room temperature. After shaking, beads were pelleted on a 96-well plate magnet and washed two times for 2 minutes with 200µL of 1x PBS-TBN. Beads were pelleted on the magnet and the wash solution removed. ACE2-biotin (100µL at 0.25µg/mL, Sino Biological Inc #: 10108-H08H-B) was added to each well. Plates were covered with foil and shaken at 700rpm for 1 hour at room temperature. After shaking, beads were pelleted on a 96-well plate magnet and washed two times for 2 minutes with 200µL of 1x PBS-TBN. Beads were pelleted on the magnet and the wash solution removed. Streptavidin-phycoerythrin (MOSS INC: SAPE-001) (100µL at 4µg/mL) was added to each well. Plates were covered with foil and shaken at 700rpm for 30 minutes at room temperature. After shaking, beads were pelleted on a 96-well plate magnet and washed two times for 2 minutes with 200µL of 1x PBS-TBN. Beads were resuspended in 100µL of 1x PBS-TBN and analyzed on a Luminex MagPix. Median fluorescent intensity values were plotted against serum dilution factor, and a sigmoidal regression line was fitted to the data using GraphPad Prism v9.0.0. Calculated half maximal inhibitory concentration (IC_50_) values of the sigmoidal curves were plotted separately as a measure of neutralizing capacity.

### Cytokine analysis of serum-treated SARS-CoV-2 VoC infected mice

Virus-inactivated serum samples or lung supernatants from SARS-CoV-2 VoC infected mice were added to a custom 8-plex Mouse Magnetic Luminex Assay (R&D Systems LXSAMSM-08) including IL-6, TNF, IFN-γ, IL-10, IL-27, IL-1β, IL-2, IL-13, and IL-17 at the recommended dilution factor (2-fold dilution). Cytokine arrays were read on a Luminex MagPix instrument.

### Flow cytometry of SARS-CoV-2 infected K18-hACE2 transgenic mouse lungs

Lung homogenates were thawed and pelleted at 1000 x *g* for 5 minutes at 4°C. PharmLyse (BD Biosciences 555899, 1mL of 1X solution) was added to each sample and homogenates were incubated for 2 minutes at 37°C. Samples were then pelleted at 1000 x *g* for 5 minutes at 4°C. Pellets were resuspended in 300µL of PBS + 1% v/v FBS and 150µL were added to 2µL of Mouse BD F_c_ Block (BD Biosciences 553142). Homogenates in F_c_ block were incubated for 15 minutes at 4°C. After blocking, 100µL were transferred into 2µL of antibody cocktail including 0.5µg of: hamster anti-mouse CD3e BV510: BD Biosciences 563024, rat anti-mouse APC-Cy7: BD Biosciences 552051, rat anti-mouse CD11b BB515: BD Biosciences 564454, rat anti-mouse CD8a BD Biosciences 551162, rat anti-mouse CD45 PE 553081, rat anti-mouse Ly6g PerCP-eFluor710: Thermo-Fisher Scientific 46-9668-82. Samples were incubated with antibody cocktail for 1 hour at 4°C in the dark. After staining, cells were pelleted at 1000 x *g* for 5 minutes at 4°C. Pellets were washed in 500µL PBS + 1% FBS (Gibco 10437028) and re-pelleted at 1000 x *g* for 5 minutes at 4°C. Cells were resuspended in 4% paraformaldehyde and fixed for 1 hour at room temperature. Fixed cells were subsequently pelleted at 1000 x *g* for 5 minutes at 4°C, filtered through at 100µm mesh filter, resuspended in PBS + 1% FBS and analyzed on a BD LSRFortessa flow cytometer.

### Statistical analyses

All statistical tests were performed on groups with n > 3 in GraphPad Prism v9.0.0. To compare two-groups, student’s *t*-tests were used. To compare three or more groups, one-way ANOVA (parametric data) or Kruskal-Wallis (non-parametric data) were used followed by Tukey’s (parametric data) or Dunn’s (non-parametric data) multiple comparisons tests. To compare grouped data, two-way ANOVA with no correction was performed followed by Tukey’s multiple comparison test. To assess statistical differences between Kaplan-Meyer curves, Mantel-Cox log-rank tests were performed.

## RESULTS

### Analysis of SARS-CoV-2 VoCs in cell culture

Viral propagation of SARS-CoV-2 VoCs relative to ancestral SARS-CoV-2 strains is not fully characterized. Vero E6 cells were infected with WA-1, B.1.1.7, and B.1.351 to investigate whether these VoCs infect cells differently using standard plaque assays and viral growth kinetics. Plaque morphology of SARS-CoV-2 VoC infected cultures was distinct (Figure 1A), with B.1.1.7 resulting in a wider and rounder plaque phenotype relative to WA-1 and B.1.351 infected cells. B.1.351 viral titer was significantly increased 24 hours post-infection in cell culture but declined to levels comparable to B.1.1.7 and WA-1 over 72 hours (Figure 1B-F). To determine the genetic background of the SARS-CoV-2 VoCs used in this study, deep-sequencing was performed, and mutations relative to the WA-1 strain were identified for B.1.1.7 (Supplementary Figure 2A) and B.1.351 (Supplementary Figure 2B) prior to use in both *in vitro* and K18-hACE2 experiments. Both strains exhibited mutations associated with widely propagating B.1.1.7 and B.1.351 SARS-CoV-2 strains (Rambaut et al., 2020; Tegally et al., 2020). In addition, authentication of B.1.351 stocks used in this study revealed a large deletion in viral ORF7a (Supplementary Figure 3) that was not previously reported.

### Clinical disease progression of mice infected with SARS-CoV-2 VoCs

Enhanced infectivity and divergent genomes of SARS-CoV-2 VoCs found in humans suggests that infection of pre-clinical animal models of SARS-CoV-2 by VoCs may be different. Many features of WA-1 SARS-CoV-2 disease progression have previously been described in this model (Hassan et al., 2020; Kumari et al., 2020; Liu et al., 2021; Moreau et al., 2020; Pandey et al., 2020; Sarkar and Guha, 2020; Silvas et al., 2021). K18-hACE2 transgenic mice were infected with SARS-CoV-2 VoCs and assessed daily until moribund (Supplementary Figure 1). Physical assessments of mice were compared using 10^3^, 10^4^, and 10^5^ pfu doses. Temperature (Figure 2A-D) and weight (Figure 2E-H) were monitored for the duration of the infection. Mice infected with the B.1.1.7 strain exhibit earlier hypothermia and weight loss relative to either the B.1.351 or WA-1 VoCs (Figure 2A-D). Hypothermia and weight loss trended higher in the B.1.1.7 VoC infected mice at both 10^4^ and 10^5^ pfu doses at five days post-infection (Figure 2A-H). Clinical scores were assigned to mice (scale described in methods and Supplementary Figure 1) based on their appearance, eye closure, respiration, activity, hypothermia, and weight loss (Supplementary Figure 1) as described previously. B.1.1.7 and B.1.351 VoC infected mice exhibited higher cumulative clinical scores than the WA-1 strain at the 10^3^ pfu dose (Figure 2I) and individual clinical scores at five days post-infection increased significantly with viral dose irrespective of the viral strain (Figure 2J). Mice infected with the B.1.1.7 VoC at 10^4^ pfu exhibited an increase in clinical score earlier than the B.1.351 and WA-1 strains at the same dose (Figure 2I-J). At later time-points, both VoC presented higher clinical scores than the WA-1 at all the challenging doses studied. Importantly, at 10^3^ pfu both VoC presented 100% mortality; however, WA-1 presented 60% survival (Figure 3, Supplementary Figure 4). Mice infected with 10^4^ and 10^5^ pfu of each VoC succumbed to infection (Figure 3). The WA-1 strain exhibited the largest difference in survival between challenge doses, with approximately 50% of mice recovering from the 10^3^ pfu dose (Figure 3, Supplementary Figure 4) compared to 0% recovery in B.1.1.7 and B.1.351 challenged groups (Figure 3). Interestingly, mice infected with the B.1.351 SARS-CoV-2 VoC succumbed to infection around 5 days post-infection independently of the challenge dose used (Figure 3). These data demonstrate differences in lethality between VoC doses in the K18-hACE2 transgenic murine host.

### VoCs escape protection from convalescent plasma

Our significant differences observed in the viral genetic sequence, viral replication *in vitro*, and infection *in vivo*, established that SARS-CoV-2 VoC behave differently during infection driving differential outcome. Given the evidence that SARS-CoV-2 VoCs can resist antibody therapeutics (Wang et al., 2021), we next sought to investigate if VoCs may bypass protection from antibodies present in convalescent plasma (CP) from an individual infected with SARS-CoV-2 in March of 2020. K18-hACE2 transgenic mice were intranasally challenged with a lethal dose of 10^5^ pfu of SARS-CoV-2 VoCs B.1.1.7, B.1.351, or WA-1 and subsequently treated with CP obtained from an individual infected in March 2020 (WA-1), or from non-SARS-CoV-2 exposed healthy human serum (HHS, confirmed by PCR and serological testing) (Figure 4). IgG levels and nAb data for these sera relative to other serum samples from SARS-CoV-2 infected, non-infected, and a vaccinated human is provided (see Supplementary Figure 5). Mice infected with the B.1.1.7 VoC and WA-1 strain exhibited significant differences in clinical measurements when treated with HHS or CP. Trends in temperature (Figure 4A-C), weight loss (Figure 4E-G). and clinical score (Figure 4I) over time were different between HHS and CP treated groups. B.1.1.7 and WA-1 exhibited significantly reduced temperature four days post-infection, B.1.1.7 exhibited significantly reduced weight four days post-infection, and B.1.1.7 and WA-1 exhibited significantly increased clinical scores four days post-infection when treated with HHS compared to CP (Figure 4). All WA-1 infected mice had no temperature or weight loss, and 60% of B.1.1.7 infected mice had no temperature or weight loss when treated with CP (Figure 4A-H). However, all mice infected with the B.1.351 VoC exhibited observable declines in temperature and weight regardless of treatment, suggesting that CP does not adequately protect mice from B.1.351 infection. Survival of mice treated with CP was significantly different than mice treated with HHS for the B.1.1.7 and WA-1 VoCs (Figure 5A-B). CP protected 100% of mice against lethal WA-1 infection and 60% of mice against lethal B.1.1.7 infection. However, no protection was observed in mice infected with the B.1.351 VoC, with 100% of CP-treated infected-mice reaching morbidity one day later (6 days post-infection) than B.1.351 infected mice treated with HHS (5 days post-infection). (Figure 5A-B). CP treatment resulted in decreased viral copy number in the lungs of mice infected with either WA-1 or B.1.1.7, but not mice infected with B.1.351 (Figure 5C). CP treatment only reduced viral copy number in brain tissue of mice infected with WA-1 and not differences in viral titers were observed in mice infected with B.1.1.7 or B.1.351 (Figure 5D).

**Figure 5.**
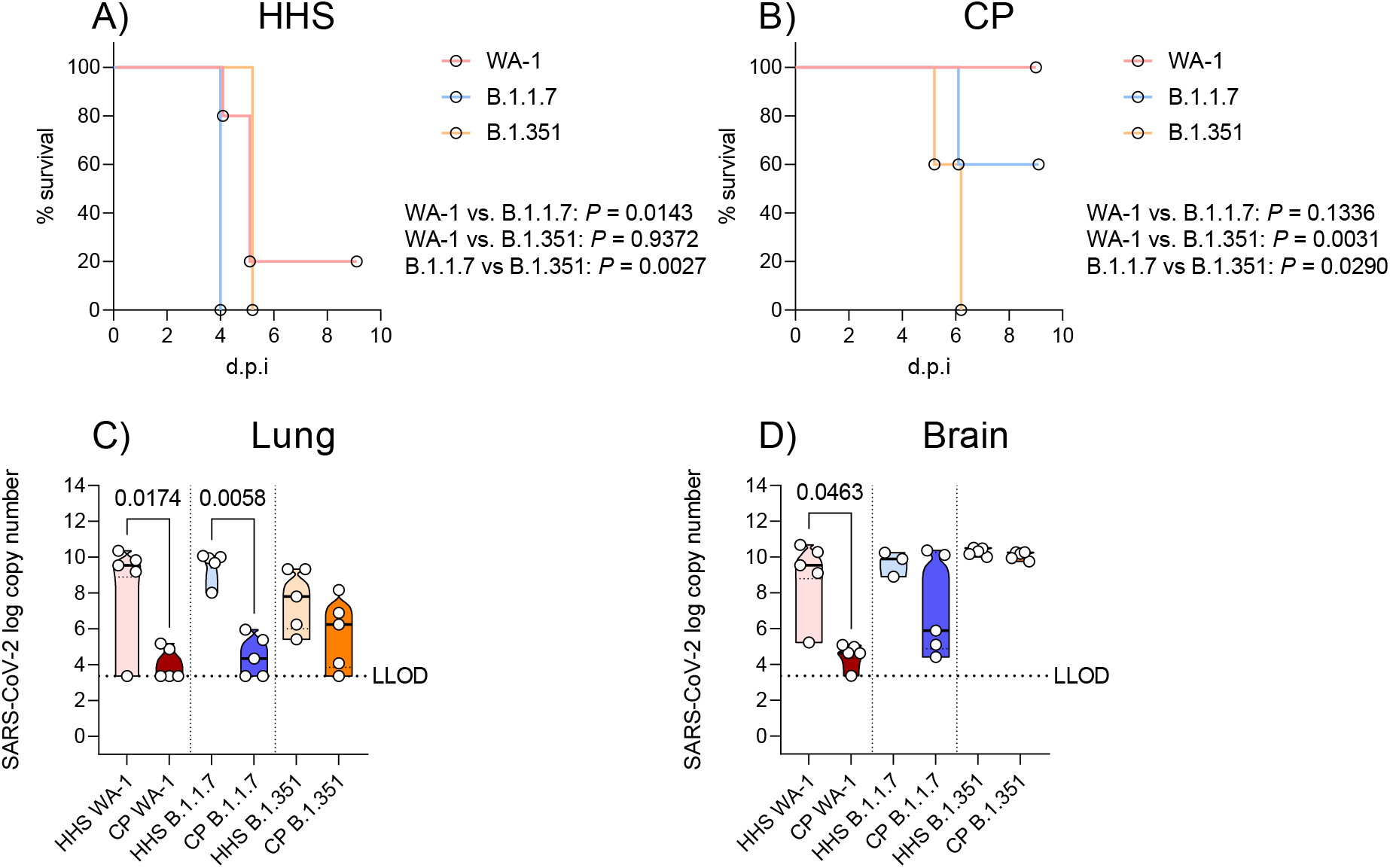
Survival and viral infection of serum-treated K18-hACE2 transgenic mice infected with SARS-CoV-2 VoCs: Kaplan-Meyer survival curves of mice infected with B.1.1.7, B.1.351, or WA-1 treated with HHS (A) or early pandemic SARS-CoV-2 CP (B). Viral copy numbers in the lung (C) and brain (D) of infected mice. LLOD = lower limit of detection based on a standard curve. Statistical significance of survival curves was assessed with the Mantel-Cox test. Statistical significance between viral copy number was assessed by a Kruskal-Wallis test followed by Dunn’s multiple comparisons test. n > 3 subjects per group. *P* values for significant differences are reported.

### Human and mouse IgG levels in convalescent plasma treated K18-hACE2 transgenic mice infected with SARS-CoV-2 VoCs

To determine the level of IgGs delivered to HHS and CP treated mice, we analyzed whether human anti-SARS-CoV-2 IgGs were present within the lung and sera of animals treated with CP or HHS through the course of infection (Figure 6, Supplementary Figure 6). The data demonstrate that significant quantities of human anti-SARS-CoV-2 IgGs targeting both the S RBD and N proteins were present at two days post-infection (Supplementary Figure 6), in sera (Figure 6A, 6C, 6E, 6G) and lung (Figure 6B, 6D, 6F, 6H) at euthanasia of CP-treated mice relative to HHS-treated mice. Relative quantities of anti-SARS-CoV-2 IgGs were similar across CP groups at all time points. A non-significant decrease of anti-N IgGs was observed in the lungs of CP-treated B.1.351 infected mice (Figure 6H). This observation suggests that there may be a greater prevalence of free N antigen in the lung that is decreasing the anti-N IgG titer. These data demonstrate successful administration of anti-SARS-CoV-2 IgGs to CP-treated mice.

**Figure 6.**
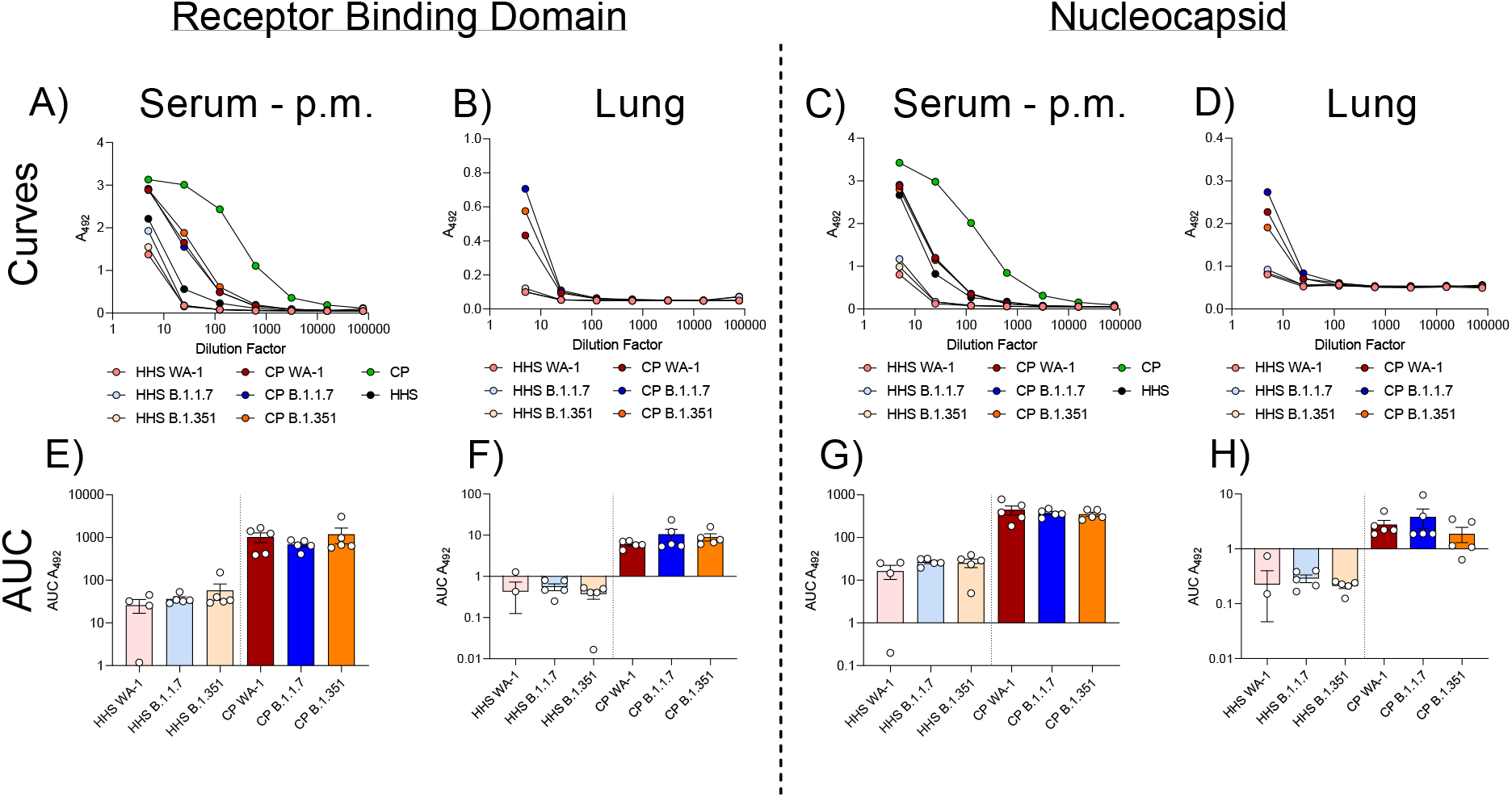
Human anti-SARS-CoV-2 IgGs in serum-treated K18-hACE2 transgenic mice infected with SARS-CoV-2 VoCs: Human antibody (IgG) levels against SARS-CoV-2 antigens. Anti-RBD curves in serum (A) and lung (B) or anti-N curves in serum (C) or lung (D) of infected mice. Area under the curve (AUC) analyses of anti-RBD IgG levels in the serum (E) or lung (F) of infected mice. AUC analyses of anti-N IgG levels in the serum (G) or lung (H). Statistical significance between AUCs was assessed by a Kruskal-Wallis test followed by Dunn’s multiple comparisons test.. n > 3 subjects per group.

### Immunological response to VoC infection and treatment with convalescent plasma

Next, we assessed cytokine levels in K18-hACE2 transgenic mice infected with SARS-CoV-2 VoCs. Serum from animals at 2 days post-infection and at euthanasia, or supernatants from lung homogenates were tested for the presence of Th1 (TNF, IL-2). Thh2 (IL-6, IL-10, IL-13), inflammasome (IL-1β) and regulatory (IL-27) cytokines (Figure 7, Supplementary Figure 7). We assayed these cytokines as many are established as pro-inflammatory mediators that are upregulated during SARS-CoV-2 infection (Hojyo et al., 2020; Horspool et al., 2021; Oladunni et al., 2020; Ye et al., 2020b) or are involved in the response to viral encephalitis (Angioni et al., 2020; Aquino et al., 2021; Fabbi et al., 2017; Oladunni et al., 2020), including K18 hACE2 transgenic mice. CP treatment reduced IL-6, TNF-α, IFN-γ, IL-10 and IL-27 in B.1.351 infected mice, and IL-6 in B.1.1.7 two days post infection (Figure 7A-E). This trend was abrogated or reversed in B.1.351 infected mice at euthanasia (Figure 7F-J). Limited differences were observed in cytokine expression in the lung, except decreased TNF-α in the lungs of CP-treated mice infected with WA-1 (Figure 7K-O). Minor or no differences were detected in IL-1β, IL-2, IL-13, and IL-17 at two days post-infection, euthanasia, or in the lung (Supplementary Figure 7). To understand the cellular response to infection, we analyzed T cells and myeloid cells in the lungs of K18-hACE2 transgenic mice infected with SARS-CoV-2 VoCs. B.1.351 infected mice exhibited significantly increased T-cell (CD3^+^ or CD4^+^) recruitment to the lungs relative to WA-1 infected mice (Supplementary Figure 8A-B). Non-significant trends in CD8^+^ T cells and myeloid cells were observed but further studies defining their function will be required (Supplementary Figure 8C-F).

**Figure 7.**
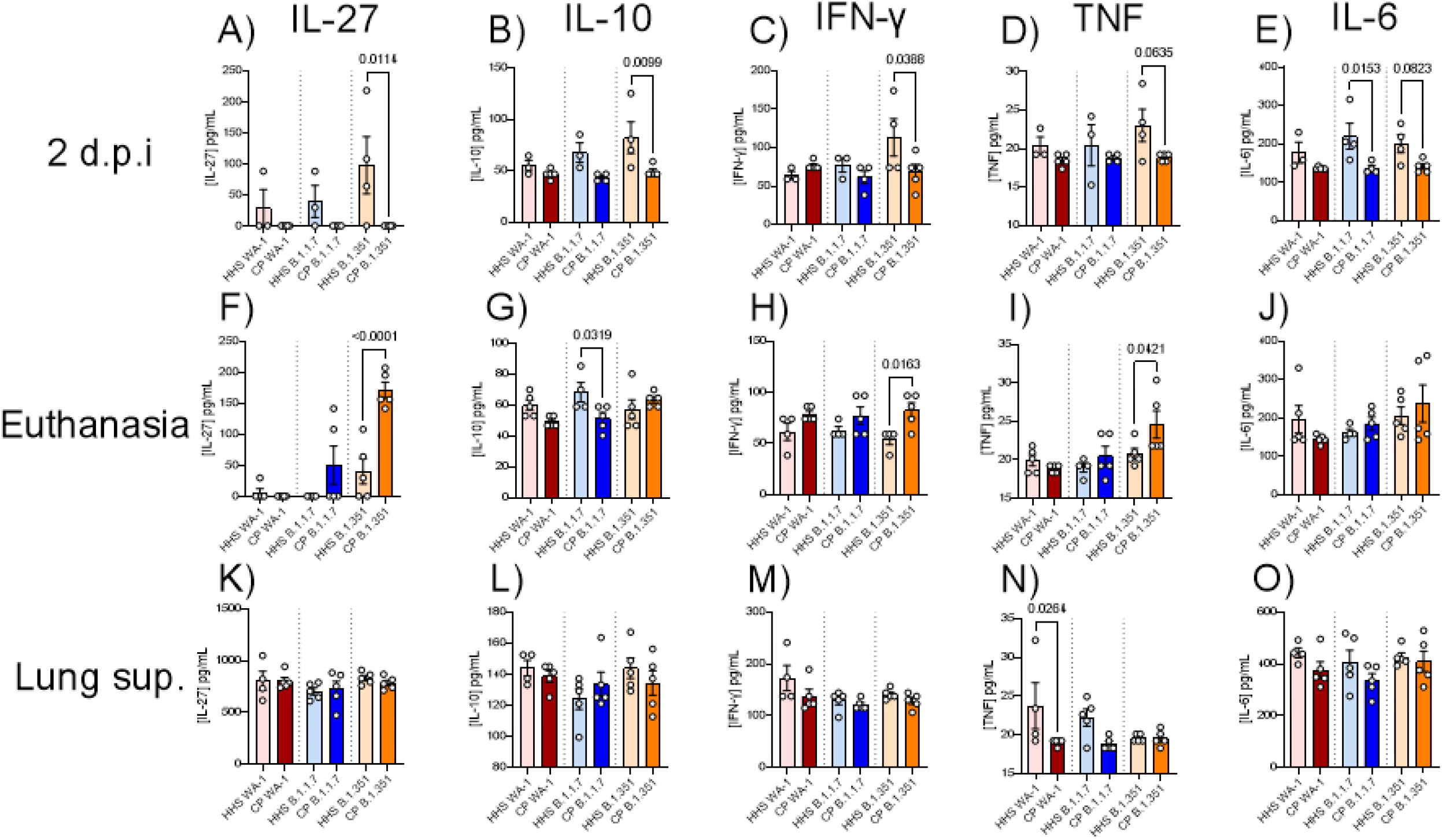
Cytokine responses in serum-treated K18-hACE2 transgenic mice: IL-6, TNF-a, IFN-y, IL-10, and IL-27 were quantified in serum two days post infection (A-E), in serum at euthanasia (F-J), or in the lung (K-O). Statistical significance was assessed by one-way ANOVA followed by Tukey’s multiple comparison test. n > 3 subjects per group. *P* values for significant differences are reported.

## DISCUSSION

SARS-CoV-2 VoCs of concern are rapidly evolving and some are exerting dramatic negative impacts on the currently ongoing COVID-19 pandemic. The goal of this study was to gain information regarding the infectivity of SARS-CoV-2 VoCs in *in vitro* and *in vivo* using the validated K18 hACE2 transgenic mouse model of SARS-CoV-2 infection. Our data present a broad picture suggesting that the VoCs exhibit major differences in pathogenesis, immune activation, and lethality in the K18-hACE2 transgenic mouse model of infection. B.1.351 replicated faster in cell culture than B.1.1.7 and WA-1. Both B.1.1.7 and B.1.351 triggered severe clinical indications of disease at lower infectious doses than the ancestral WA-1 strain in the K18-hACE2 transgenic murine model. Treatment of infected mice with CP from WA-1-infected individuals (March of 2020, prior to the emergence of B.1.1.7 and B.1.351 VoC) resulted in full protection only against the WA-1; B.1.1.7 infected mice exhibited partial protection and B.1.351 exhibited no protection. Both VoCs resulted in significant viral replication in the brain of K18-hACE2 transgenic mice despite treatment with CP. It is clear by comparing all results from this study (Supplementary Figure 9) that CP efficacy is significantly reduced against SARS-CoV-2 VoC and that antibodies against ancestral SARS-CoV-2 may not be fully protective against B.1.1.7 and B.1.351. This may have additional impacts on the efficacy of existing SARS-CoV-2 vaccines that target the S protein of WA-1 SARS-CoV-2. Ultimately, these results provide a troubling picture of the impact of VoCs on SARS-CoV-2 pathogenesis and host immunity.

SARS-CoV-2 B.1.351 infection resulted in unique phenotypes in this study: increased replication in cell culture, clinical presentations at low doses of infection, lethality at low infectious doses, reduced clinical and survival benefits from early (WA-1) CP, increased viral load in lung and brain tissues, and increased pro-inflammatory cytokines after CP treatment, all supporting the conclusion that the B.1.351 VoC results in more severe disease than the original WA-1 strain in this model. Of particular concern is the lack of reduction in viral copy number in all mice treated with CP infected with B.1.351 relative to WA-1 infected mice. In addition, the severity of viral burden despite CP treatment was enhanced in B.1.351 infected mice relative to B.1.1.7 infected mice: 100% of CP-treated B.1.351 infected mice had lung and brain infection relative to 40% of CP-treated B.1.1.7 infected mice. This demonstrates that B.1.351 infection may harbor more resistance or tolerance to early pandemic CP than B.1.1.7. It is important to point out the caveat that we did not investigate antibody dependent enhancement of infection. The observed differences in our mouse challenge studies may be attributable to the alterations in the viral B.1.351 S RBD region of the S protein, which may increase affinity for the hACE2 receptor (Bozdaganyan et al., 2021; Laffeber et al., 2021; Ozono et al., 2021; Shah et al., 2020; Tian et al., 2021) and could result in increased infectivity and pathogenicity. Although this yet speculative, sequencing results in Supplementary Figure 2-3 illustrate that many other mutations exist within the B.1.351 SARS-CoV-2 genome. Mutations within other viral proteins, particularly those involved in viral replication and genome packaging may provide additional advantages for this VoC during infection. It is currently unknown what impact the B.1.351 ORF7a deletion we observed has on pathogenesis, but other studies have suggested limited impacts of similar ORF7a mutations in cell culture (Chiem et al., 2020; Narayanan et al., 2008; Sims et al., 2005). Dissecting the consequences of these B.1.351 non-S mutations will be useful in determining the mechanisms behind the increased disease severity observed here and may help better inform containment of these VoCs, and others emerging in the future.

Several additional observations were made in this study, including discovering significant differences in cytokine production of B.1.351 SARS-CoV-2 VoC infected mice. IL-27 is an indicator of acute encephalitis and can activate CD8+ T cells (Angioni et al., 2020; Aquino et al., 2021; Fabbi et al., 2017) in addition to several other less characterized functions (Awasthi et al., 2007; Iwasaki et al., 2013; Jung and Robinson, 2014; Kastelein et al., 2007; Seman et al., 2020; Sugiyama et al., 2008). IL-27 also regulates IL-10 production, the latter acting in an anti-inflammatory manner by suppressing inflammatory responses (Iyer and Cheng, 2012). IL-27 was increased in B.1.351 infected mice. B.1.351 challenged mice also had significant viral burden in the brain, increased CD3^+^ T cells in the lung, and non-significantly increased in CD8+ T cells in the lung. The combination of these data suggests that IL-27 expression in B.1.351 infected mice may be linked to more severe encephalitis and potentially higher activation of CD8^+^ T cells. HHS-treated mice infected with B.1.351 exhibited this increase in IL-27 at two days post-infection, whereas CP-treated mice exhibited this increase only at euthanasia. It is possible that this delay in increased IL-27 may be beneficial as CP-treated B.1.351 infected mice survive for one additional day over HHS-treated mice. Mechanistically, this may be due to IL-27 promoting uncontrolled inflammation (Takeda et al., 2003) and suppression of T_regs_ (Cox et al., 2011) which may increase local and systemic damage. However, IL-27 could also be increased as an emergency response in response to encephalitic infection and may exert beneficial pro- or anti-inflammatory effects dependent on the severity of infection. Further studies will be required to elucidate the true role of IL-27 in this system. In a similar manner, IL-6, TNF, and IFN-γ were increased in B.1.351 infected mice two days post-infection in HHS-treated mice and increased in CP-treated mice at euthanasia. This delay in induction of pro-inflammatory cytokines may be beneficial, but ultimately none of these delays appear to protect mice from death.

This study also provides additional insights on VoC pathogenicity; however, several important clarifications must be made when interpreting these results. Firstly, although these infection models exhibit clear differences in infectivity, infections in cell culture and mice do not perfectly represent those in humans. In particular, the K18-hACE2 transgenic mouse model is a lethal disease model, with different viral tropism, which frequently can result in 100% mortality at low infectious doses as seen in this study and others (Hassan et al., 2020; Kumari et al., 2020; Liu et al., 2021; Moreau et al., 2020; Oladunni et al., 2020; Pandey et al., 2020; Sarkar and Guha, 2020; Silvas et al., 2021). Human disease can range from mild disease to lethal disease and is impacted by comorbidities (Elezkurtaj et al., 2021; Mehra et al., 2020; Wu and McGoogan, 2020). Re-creating this in any animal model is exceptionally challenging. Another complex component is that it is unknown what the minimum infectious dose of SARS-CoV-2 is for humans, and thus it is difficult to compare animal models of infection to a real human infection without this basic information. Secondly, the K18-hACE2 transgenic mouse expresses hACE2 but also retains expression of murine (m)ACE2. There is speculation that SARS-CoV-2 VoCs can infect wild-type mice and cause non-lethal disease by binding to mACE2 (Montagutelli et al., 2021); this may impact the comparisons provided in this study. We have observed similar phenomena in preliminary studies where some of these VoC are able to infect wild-type mice, contrary to the situation with WA-1 strain (data not shown). Although this may be true, infection of Vero E6 cells with B.1.351 (a non-human cell line that is not genetically modified for infection) was significantly increased 24 hours post-infection. This provides support of the conclusions presented here in light of the potential for VoCs to bind mACE2, although an effect of mACE2 cannot be excluded. CP as a treatment has been used widely since the onset of the COVID-19 pandemic (Bloch, 2020; Bloch et al., 2020; Chen et al., 2020), but its efficacy has recently been re-evaluated under certain conditions (Casadevall et al., 2021; Cele et al., 2021; NIH, 2021; Zhao and He, 2020). It is possible that some of the decreased efficacy of CP in this study (and others) may be due to SARS-CoV-2 VoCs, and the data presented here support this theory. Other factors including lack of adequate screening for nAbs and lot-to-lot variability of plasma samples may be the cause of these differences in larger studies of humans treated with CP. The plasma used in this study was determined to have high levels of IgG antibodies against SARS-CoV-2 S RBD and N proteins and also exhibited neutralizing activity similar to a vaccinated individual (Supplementary Figure 4). In this context, this plasma should have high protective capacity, and indeed protected WA-1 infected mice against severe disease. Consequently, it is concerning that protection is limited or absent when K18-hACE2 transgenic mice were challenged with B.1.1.7 or B.1.351 VoCs, respectively. It is important to note that only one CP sample was tested in this study and was obtained from a naturally infected, not a vaccinated, individual. Therefore, although there appears to be a lack of protective capacity of this plasma against B.1.351 and partial protective capacity against B.1.1.7, these data do not provide direct against the efficacy of SARS-CoV-2 vaccines which stimulate broader immune responses. However, it is likely that the production of antibodies elicited by existing SARS-CoV-2 vaccines are not fully protective against VoCs, which warrants caution. Further analyses of these vaccines will be critical in determining their efficacy against B.1.1.7, B.1.351, new VoCs, or variants of high concern. Continued support for novel vaccine development against VoCs will be instrumental in providing full protection against evolving strains of SARS-CoV-2.

To summarize, this study provides early insight into differences in SARS-CoV-2 VoC pathogenicity in K18-hACE2 transgenic mice and the efficacy of CP containing antibodies targeting WA-1 SARS-CoV-2 against newly emerged SARS-CoV-2 VoCs in the K18-hACE2 transgenic mouse model. This study demonstrates increased disease pathology for mice infected with B.1.1.7 and B.1.351 VoCs and lack or limited protection from CP in mice infected with B.1.351 and B.1.1.7, respectively. Immunological profiles were different between mice infected with each VoC, with B.1.351 stimulating greater increases in cytokine levels and potential T-cell recruitment, most likely due to its more efficient replication as compared to the WA-1 and B.1.1.7 strains. Overall, these data present a concerning picture of the SARS-CoV-2 VoCs warranting continued caution as the pandemic continues and evolves and suggest the need to update current vaccines to protect against these newly emerged SARS-CoV-2 VoC.

## ACKNOWLEDGEMENTS

We would like to express our gratitude to Laura Gibson and Clay Marsh for enabling this research through resources and support. This project was supported by the Vaccine Development Center at the West Virginia University Health Sciences Center. F.H.D. and the VDC are supported by the Research Challenge Grant no. HEPC.dsr.18.6 from the Division of Science and Research, WV Higher Education Policy Commission. Flow cytometry analyses were supported financially by the West Virginia University Flow Cytometry & Single Cell Core Facility, which is supported by the National Institutes of Health equipment grant number S10OD016165 and the Institutional Development Awards (IDeA) from the National Institute of General Medical Sciences of the National Institutes of Health under grant numbers P30GM121322 (TME CoBRE) and P20GM103434 (INBRE).

## AUTHOR CONTRIBUTIONS

A.M.H, F.H.D, I.M, J.T, C.Y and L.M.S designed the experiments. C.Y performed *in vitro* analyses of SARS-CoV-2 VoCs, and J.S., A.G, and R.K.P performed sequencing. M.T.W and I.M propagated virus for animal experiments. A.M.H, T.Y.W, B.P.R, K.S.L and F.H.D infected and euthanized animals, dissected organs, and prepared them for analyses. A.M.H and B.P.R performed ELISA against SARS-CoV-2 antigens and A.M.H and T.Y.W performed qPCR to determine viral load. A.M.H performed cytokine analyses. A.M.H and T.Y.W prepared cells for flow cytometry and A.M.H performed flow cytometry and analysis. J.D analyzed sequence data from viral passages used for infection of K18-hACE2 transgenic mice. A.M.H analyzed, formatted, and represented data for publication. H.A.C assisted with general revisions. All authors contributed to the writing and revision of the manuscript.

## SUPPLEMENTARY FIGURES

**Supplementary Figure 1.**
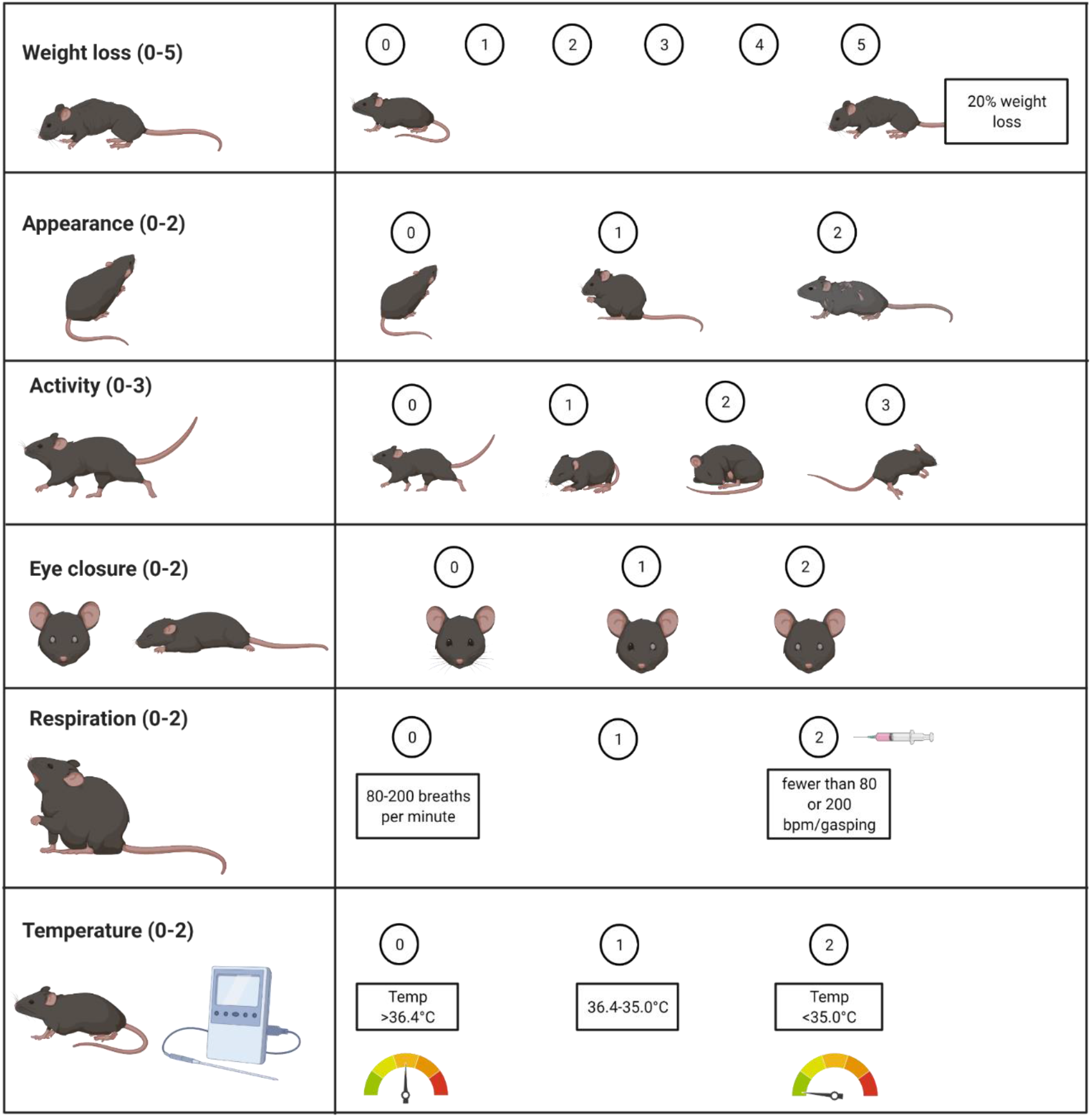
Clinical scoring system: K18-hACE2 transgenic mice were assessed for weight loss, appearance, activity, eye closure, respiration, and temperature and scored using the metrics provided. Mice with a score of 5 in weight loss or a 2 in respiration were euthanized.

**Supplementary Figure 2.**
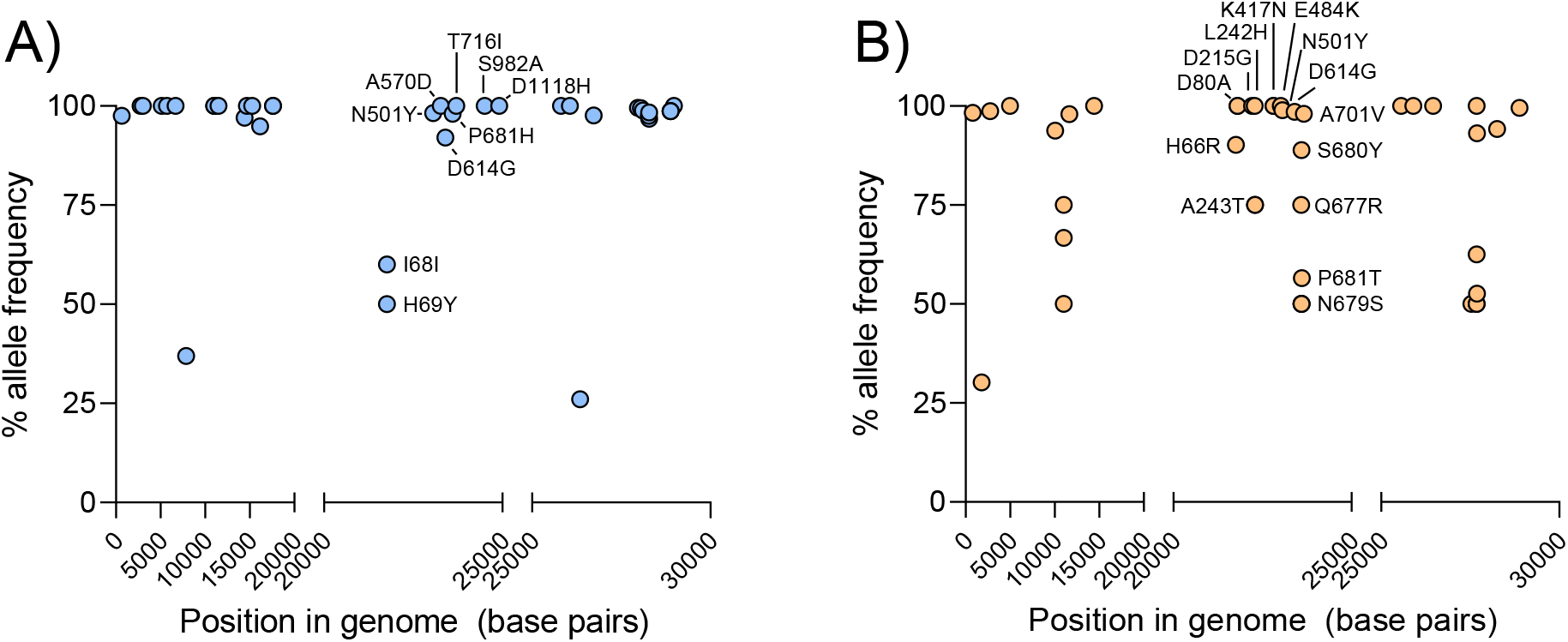
Sequencing of SARS-CoV-2 VoCs in this study: NGS of SARS-CoV-2 WT and VoCs demonstrates that mutations associated with each variant are present within the B.1.1.7 (A) and B.1.351 (B) genomes. Mutations within the spike protein are labeled.

**Supplementary Figure 3.**
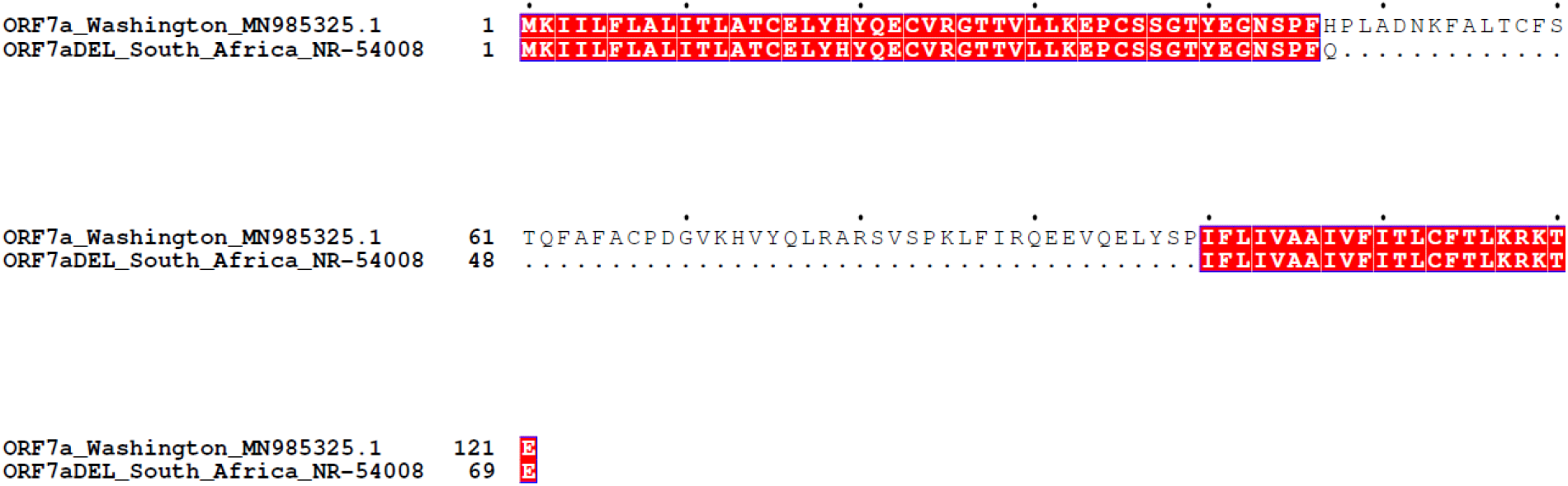
ORF7a deletion in B.1.351 SARS-CoV-2 VoC in this study: Annotated deletion within the ORF7a protein of the B.1.351 VoC used in this study.

**Supplementary Figure 4.**
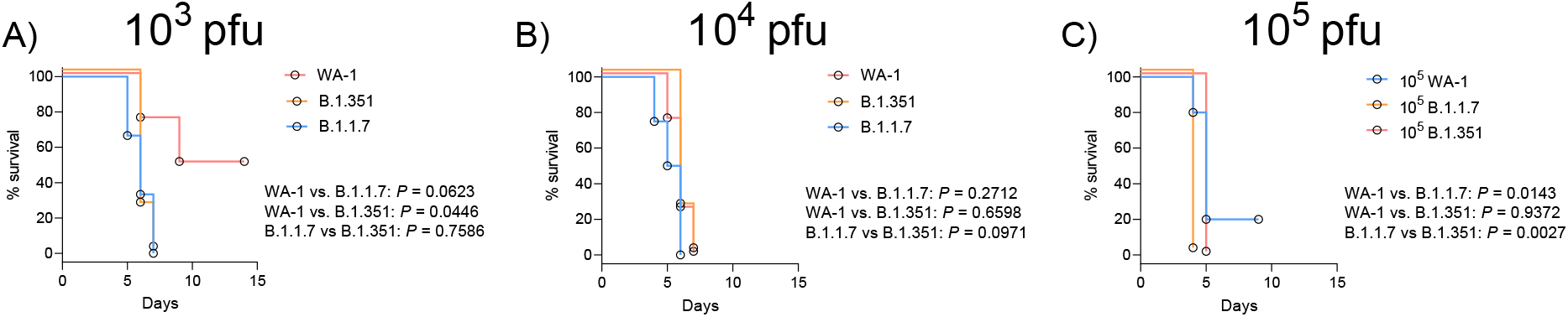
Impact of variant on survival of K18-hACE2 transgenic mice infected with SARS-CoV-2 VoCs: Kaplan-Meyer survival curves of mice infected with 10^3^ (A), 10^4^ (B), or 10^5^ (C) pfu doses of WA-1 B.1.1.7, or B.1.351 SARS-CoV-2 VoCs. Statistical significance was assessed by Mantel-Cox tests. n > 3 subjects per group. *P* values for significant differences are reported.

**Supplementary Figure 5.**
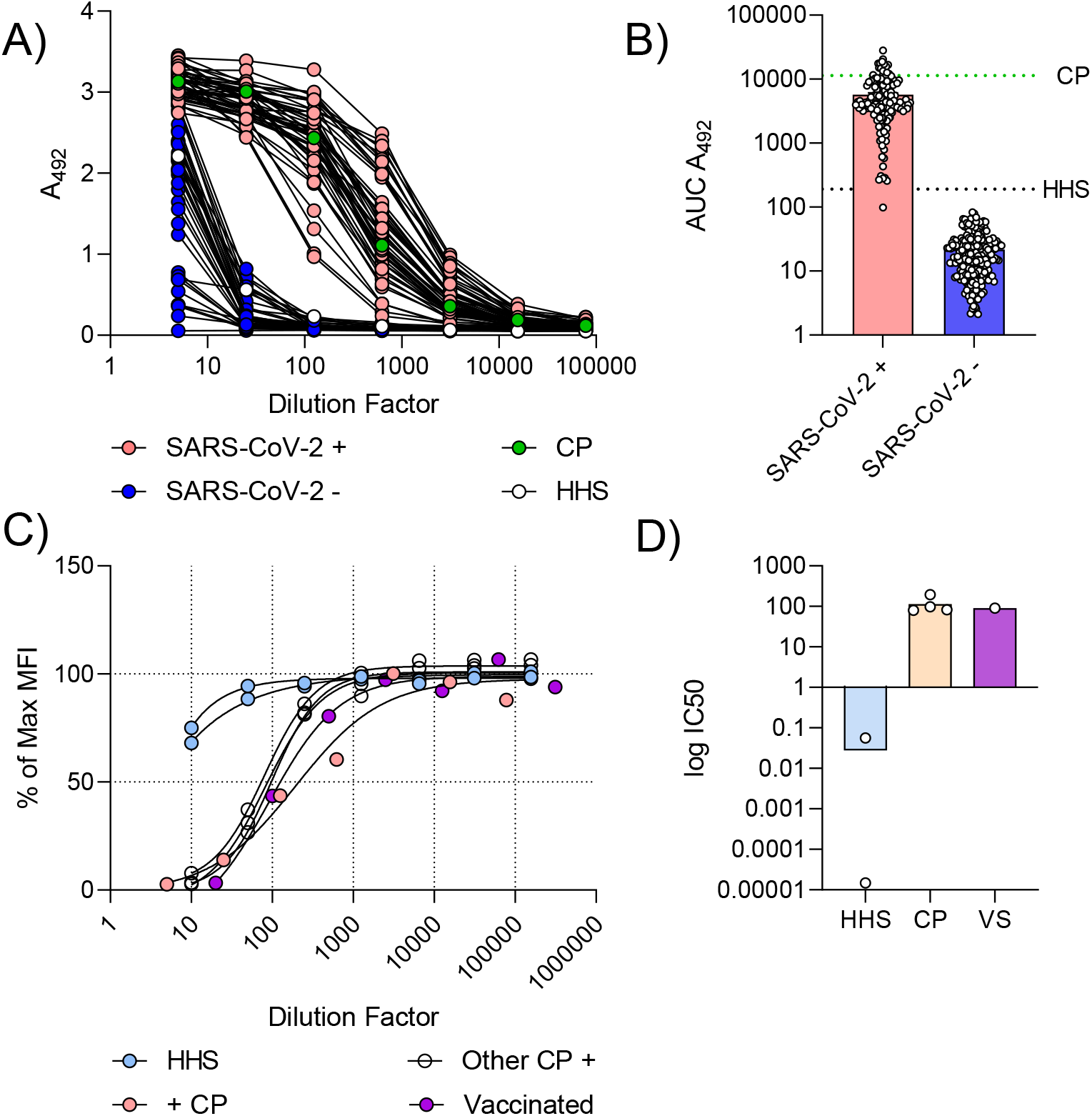
Anti-RBD IgG levels and neutralization capacity of HHS and CP: IgG levels of CP and HHS were assessed using ELISA developed previously. CP (green, dotted green line) and HHS (white, dotted black line) ELISA curves (A) and AUCs (B) were compared to a subset of SARS-CoV-2^+^ (red) and SARS-CoV-2^-^ (blue) individuals from a prior study. Neutralizing antibody function was assessed with a modified Luminex assay. Neutralization curves (C) and IC50 values (D) for nAbs from HHS (blue), CP (red), and a vaccinated individual (purple) are provided.

**Supplementary Figure 6.**
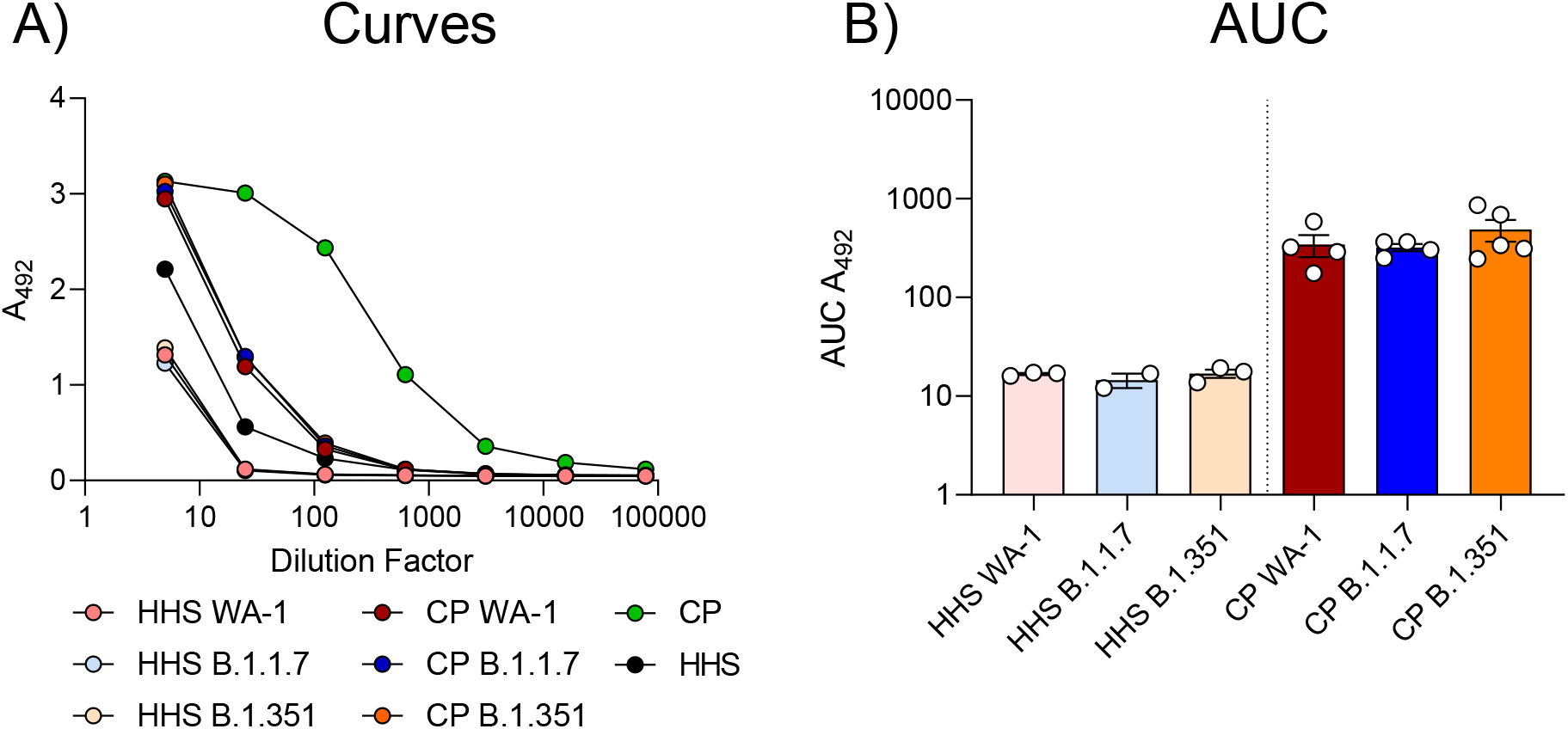
Human anti-SARS-CoV-2 IgGs two days post-infection in serum-treated K18-hACE2 transgenic mice infected with SARS-CoV-2 VoCs: Antibody (IgG) levels against SARS-CoV-2 antigens. Anti-RBD curves in serum (A) and of infected K18-hACE2 transgenic mice 2 days post-infection. Area under the curve (AUC) analyses of anti-RBD IgG levels (B). Statistical significance between AUCs was assessed by a Kruskal-Wallis test followed by Dunn’s multiple comparisons test. n > 3 subjects per group.

**Supplementary Figure 7.**
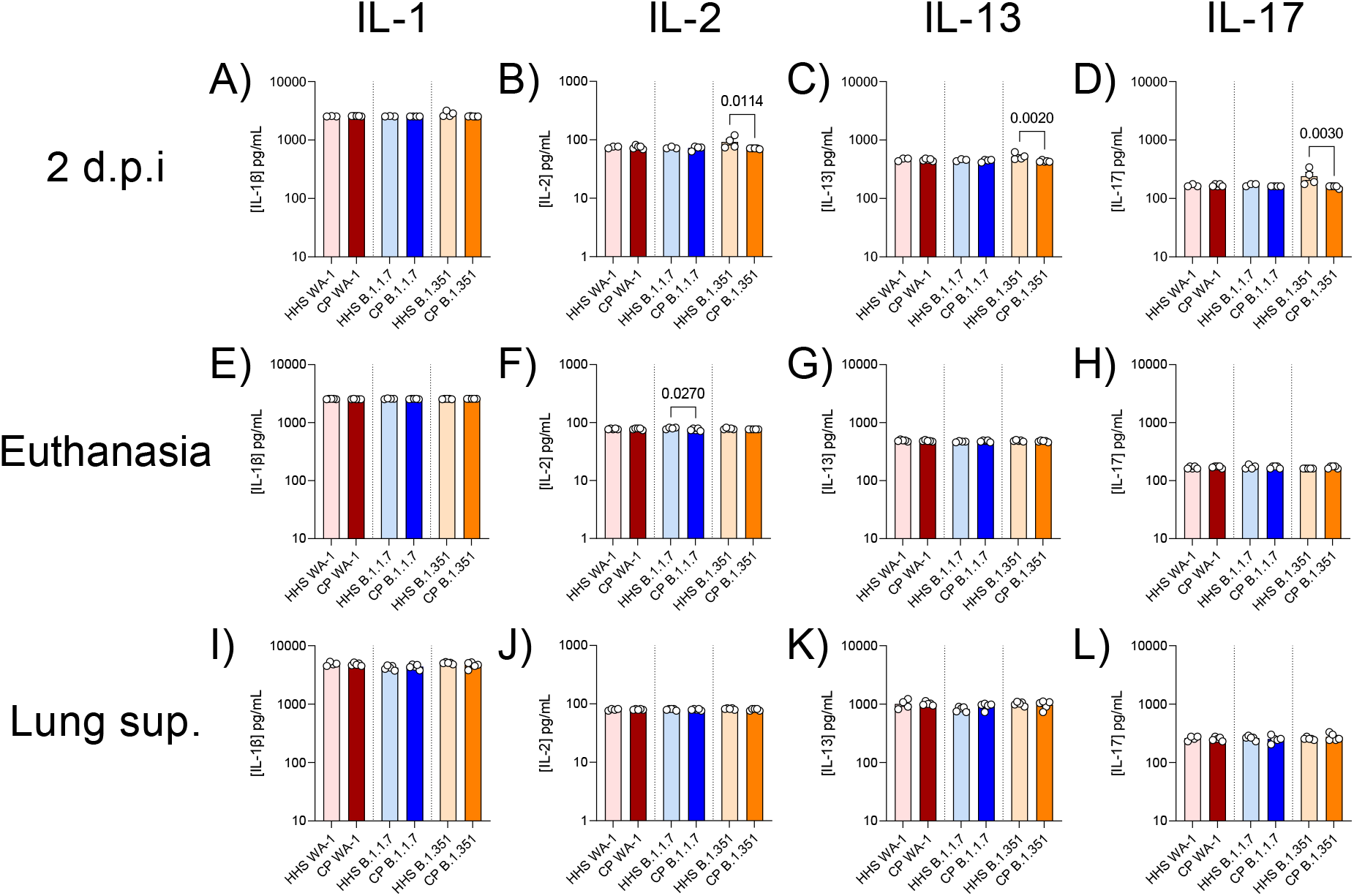
Minor or no differences in some cytokine levels in K18-hACE2 transgenic mice treated with serum: IL-1, IL-2, IL-13, and IL-17 were quantified in serum two days post infection (A-D), in serum at euthanasia (E-H), or in the lung (I-L). Statistical significance was assessed by one-way ANOVA followed by Tukey’s multiple comparison test. n > 3 subjects per group. *P* values for significant differences are reported.

**Supplementary Figure 8.**
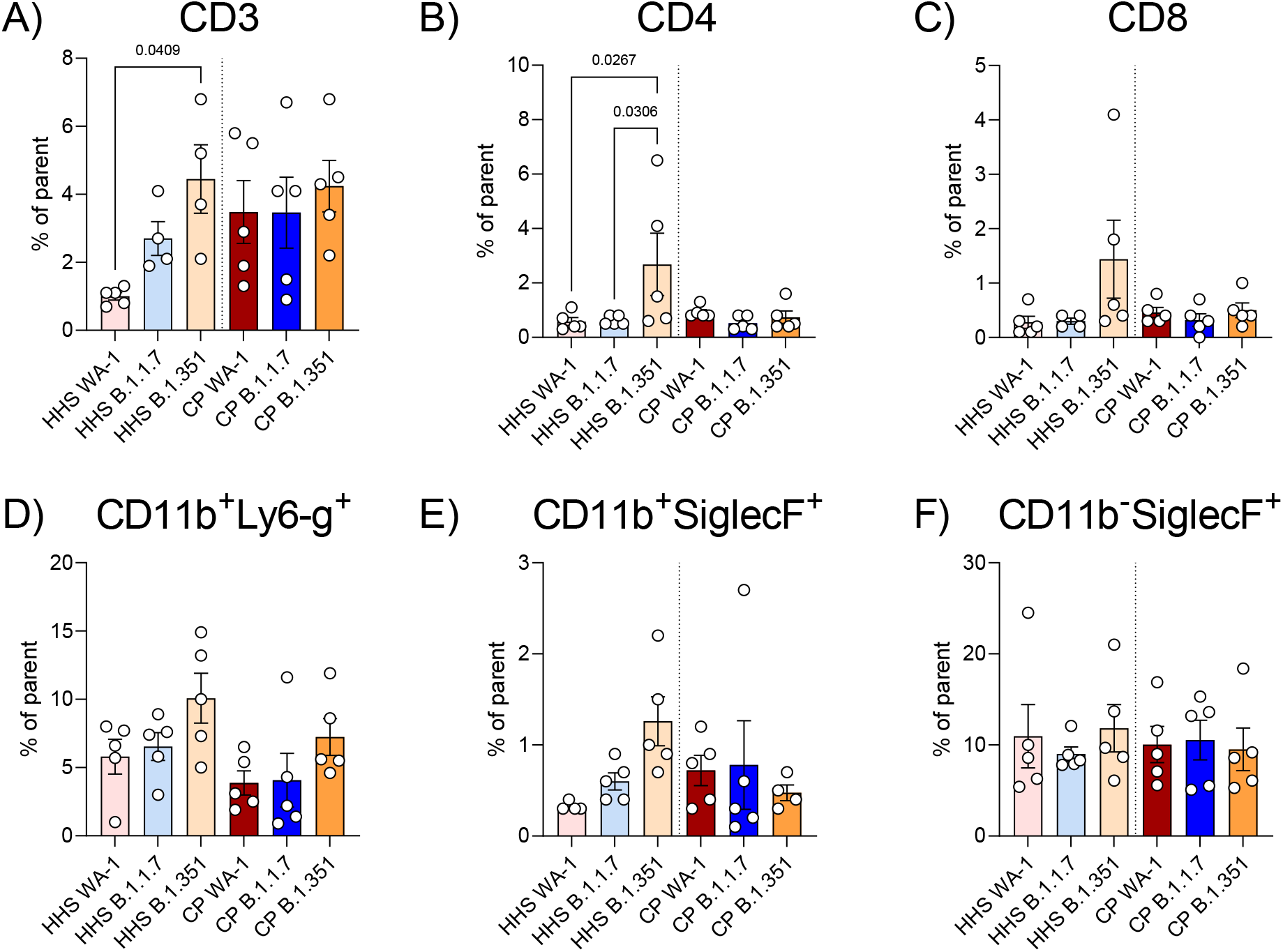
T cell and myeloid cell populations in lungs of K18-hACE2 transgenic mice infected with SARS-CoV-2 VoCs: CD3 (A), CD4 (B), and CD8 (C) T cells were quantified in lung homogenates from SARS-CoV-2 variant infected K18-hACE2 transgenic mice. CD11b^+^Ly6-g^+^ (D), CD11b^+^SiglecF^+^, and CD11b^-^SiglecF^+^ cells myeloid cells were also quantified in lung homogenates. Statistical analyses were performed by one-way ANOVA followed by Tukey’s multiple comparison test. n>3 samples per group. *P* values of statistically significant results are reported.

**Supplementary Figure 9.**
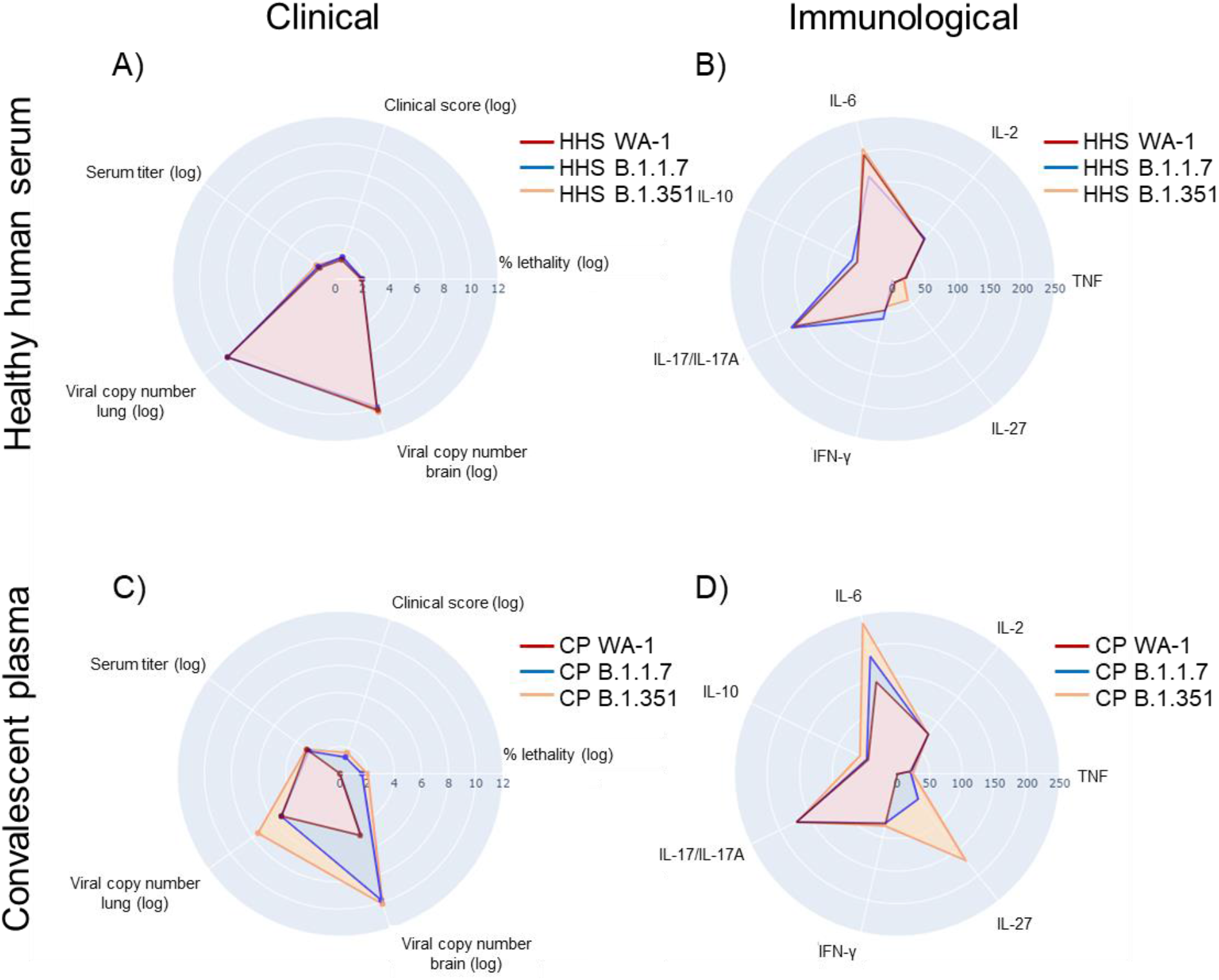
Comparison of clinical and immunological phenotypes of SARS-CoV-2 infected K18-hACE2 transgenic mice treated with HHS or CP: Clinical (A) and immunological (B) phenotypes of SARS-CoV-2 infected K18-hACE2 transgenic mice treated with HHS. Clinical (C) and immunological (D) phenotypes of SARS-CoV-2 infected K18-hACE2 transgenic mice. Scatterpolar plots were generated in Python using plotly.

